# Mapping the Nonhistone Interactome of the TRIM24 Bromodomain Using Genetically Encoded Complementary Photocrosslinkers

**DOI:** 10.64898/2026.09.09.750415

**Authors:** Ishita Bardhan, Soumyajit Chakraborty, Navya Sree Ponnaganti, Babu Sudhamalla

## Abstract

TRIM24 is a tandem epigenetic reader containing plant homeodomain (PHD) and bromodomain modules in its C-terminal region. The PHD domain recognizes unmodified lysine 4 of histone H3, whereas the bromodomain recognizes acetylated histones H3 and H4. However, whether the TRIM24 bromodomain also engages acetylated nonhistone proteins remains poorly understood. To address this question, we developed site-specific photocrosslinking probes for the TRIM24 bromodomain using interaction-based protein profiling (IBPP). 4-azido-L-phenylalanine (AzF) and 4-benzoyl-L-phenylalanine (BzF) were genetically incorporated at L922, a residue positioned within the acetyllysine-binding pocket. Both variants preserved recognition of acetylated histone peptides and underwent efficient UV-dependent photocrosslinking. Quantitative proteomics of photocaptured proteins revealed distinct profiles for the two probes, identifying NPM1 among the L922AzF-enriched proteins and CDK9 among the L922BzF-enriched proteins. Additional candidate proteins included MAP4, RAN, and SND1. ChIP-seq analysis showed overlapping genomic enrichment of TRIM24, CDK9, and NPM1 near transcription start sites of cancer-associated genes. KEGG analysis further placed the identified proteins in several signalling pathways. Collectively, this study demonstrates that site-specific photocrosslinking can expand interactome analysis of epigenetic reader domains and provides candidate nonhistone proteins associated with the TRIM24 bromodomain for further investigation.

## Introduction

Eukaryotic gene transcription is regulated by dynamic interactions between chromatin, transcription factors and chromatin-associated regulatory proteins that collectively control the accessibility and activity of genomic regulatory elements (1,2). The organization and functional state of chromatin are influenced by nucleosome positioning, histone posttranslational modifications (PTMs) and the recruitment of regulatory proteins to specific genomic regions (2,3). Histone PTMs, including acetylation and methylation, provide molecular signals that are recognized by specialized protein domains, thereby linking chromatin states to transcriptional regulation and other chromatin-dependent processes (3,4). Proteins containing such domains, commonly referred to as epigenetic reader proteins, interpret these modifications and contribute to the recruitment and regulation of transcriptional and chromatin-remodeling machinery (3,4).

Bromodomains are well-characterized epigenetic reader domains that recognize ε-N-acetyllysine residues and contribute to the recruitment of transcriptional and chromatin-associated regulatory proteins (5–7). Although bromodomains are best known for recognizing acetylated histone tails, bromodomain-containing proteins can also recognize acetylated nonhistone proteins, suggesting that their interaction landscapes may extend beyond canonical chromatin recognition (6,8). Such interactions can contribute to transcriptional regulation and other cellular processes by linking bromodomain-containing proteins to additional regulatory proteins. Consequently, defining both histone and nonhistone interactions of bromodomain-containing proteins is important for understanding their molecular functions and cellular activities.

TRIM24 (tripartite motif-containing 24), also known as transcription intermediary factor 1α (TIF1α), is a multidomain member of the TRIM family with established roles in chromatin recognition, transcriptional regulation and protein homeostasis (9,10). TRIM24 contains an N-terminal RING-B-box-coiled-coil (RBCC) region and a C-terminal plant homeodomain (PHD)-bromodomain module (Figure 1A) (11). Through interactions with nuclear receptors, including oestrogen and androgen receptors, TRIM24 contributes to hormone-responsive transcriptional programmes and has been implicated in cancer-associated transcriptional regulation (11–13). In breast cancer, TRIM24 recognizes a non-canonical histone signature and contributes to oestrogen receptor-dependent transcription, whereas in prostate cancer it has been implicated in androgen receptor-dependent transcription and tumour cell proliferation (11–13). The RING domain of TRIM24 also confers E3 ubiquitin ligase activity, and TRIM24 has been reported to regulate p53 stability (14,15). Genetic studies have further identified context-dependent tumour-suppressive functions of TRIM24, including effects on retinoic acid receptor signalling (16). Thus, TRIM24 can influence cellular processes through multiple molecular activities.

**Figure 1.**
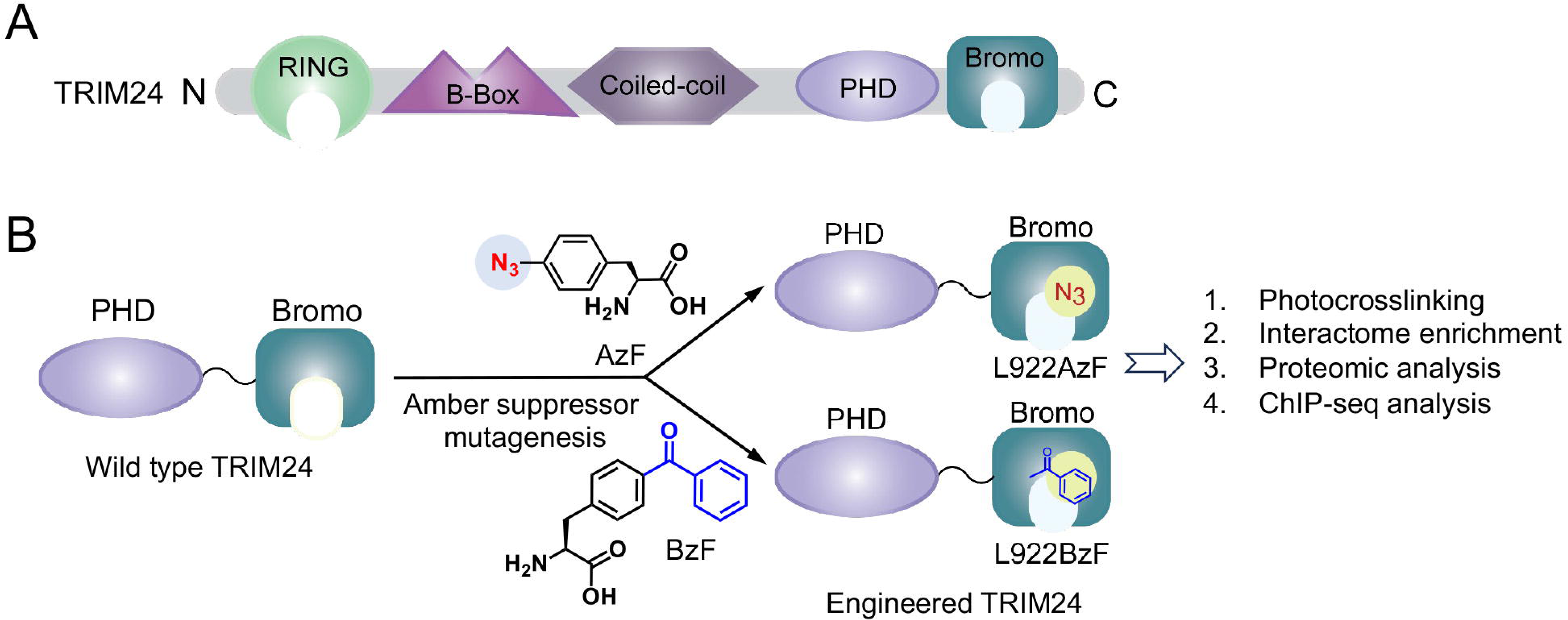
Domain organization of TRIM24 and photocrosslinking strategy for profiling its nonhistone interactome. (A) Schematic representation of the domain organization of TRIM24, comprising the N-terminal RING, B-box, and coiled-coil domains and the C-terminal PHD and bromodomains. (B) Schematic workflow for engineering the TRIM24 bromodomain with genetically encoded photocrosslinkable amino acids. An amber codon (TAG) was introduced at L922, enabling incorporation of 4-azido-L-phenylalanine (AzF) or 4-benzoyl-L-phenylalanine (BzF) by amber-suppression mutagenesis to generate TRIM24-L922AzF and TRIM24-L922BzF, respectively. Upon photoactivation, the engineered bromodomain forms covalent crosslinks with proximal proteins, enabling enrichment and proteomic identification of TRIM24-associated non-histone proteins. The identified interactome was subsequently integrated with ChIP-seq analysis to investigate the chromatin-associated functions of TRIM24.

The C-terminal PHD-bromodomain module of TRIM24 functions as an integrated epigenetic reader that enables combinatorial recognition of histone PTMs. Structural and biochemical studies demonstrated that the TRIM24 PHD-bromodomain module recognises a combinatorial histone H3 signature comprising unmodified H3K4 (H3K4me0) and acetylated H3K23 (H3K23ac), with the PHD finger and bromodomain contributing cooperatively to chromatin recognition (11). The TRIM24 bromodomain can also accommodate different acetylated histone H4 substrates. Our previous study demonstrated that the TRIM24 PHD– bromodomain module preferentially recognises H4K5ac, H4K8ac and H4K5acK8ac, while retaining recognition of H3K4me0 by the PHD finger (17). These findings establish the ability of the TRIM24 bromodomain to recognize diverse acetylated histone ligands and raise the possibility that its acetyllysine-recognition capacity may also extend to nonhistone proteins.

A precedent for such interactions is provided by BRD4, whose second bromodomain recognises acetylated TWIST with relatively weak affinity, in the order of 800 µM (6). Despite this weak interaction, BRD4 recruitment to acetylated TWIST is functionally important and, together with recognition of acetylated histone H4 by its first bromodomain, contributes to the regulation of TWIST-dependent gene expression in breast cancer cells (6). More broadly, bromodomains exhibit a range of affinities toward acetylated proteins (5,18), while conventional enrichment methods tend to favor the strongest and most abundant interactions and lack the temporal control needed to capture transient protein interactions (19). These considerations highlight the need for approaches that can covalently capture proteins in close proximity to a defined site within a bromodomain, thereby enabling the detection of weak or transient interactions that may otherwise escape conventional enrichment strategies.

Interaction-based protein profiling (IBPP) was developed to exploit this principle through the site-specific incorporation of photoreactive unnatural amino acids into reader domains (20). The initial implementation employed 4-azido-L-phenylalanine (AzF) in the BRD4 bromodomain and enabled the identification of acetylated interacting proteins (20). This strategy was subsequently extended to several bromodomain-containing proteins for interactome profiling (21,22). Our previous studies further established that both AzF and 4-benzoyl-L-phenylalanine (BzF) can be incorporated into bromodomains for site-specific photocrosslinking and interaction profiling (23). Upon irradiation, AzF generates a reactive nitrene intermediate, whereas BzF undergoes photoexcitation of its benzophenone group to generate a reactive triplet-state species capable of covalent capture of proximal proteins (24). These distinct photochemical properties provide complementary approaches for profiling bromodomain-mediated protein interactions (23,24).

Building on these findings, we sought to extend this approach to the TRIM24 bromodomain to define its acetylated protein interaction landscape. AzF and BzF were incorporated at L922, a residue positioned within the acetyllysine-binding pocket of the TRIM24 bromodomain (Figure 1B). The engineered variants were characterized for their recognition of acetylated histone ligands and photocrosslinking properties, followed by structural and molecular dynamics analyses to examine the basis of their interactions with acetylated histone peptides. We then applied cellular proteome photocrosslinking coupled with quantitative mass spectrometry to identify candidate proteins captured by the engineered TRIM24 bromodomain. Together, these approaches enabled us to investigate the acetylated protein interaction landscape of the TRIM24 bromodomain.

## Results and Discussion

### Engineering and functional characterization of TRIM24 bromodomain variants

To engineer the TRIM24 bromodomain with the unnatural amino acids AzF and BzF, we focused on the conserved “LAF shelf” motif of the TRIM24 bromodomain. Previous studies have shown that the tryptophan residue within the “WPF shelf” of the bromodomain and extra-terminal (BET) family, as well as the corresponding tryptophan-equivalent residues in non-BET bromodomains, can be substituted with unnatural amino acids without substantially affecting ligand binding (21,23). More recently, we demonstrated that the asparagine residue within the NIF shelf of the ATAD2 bromodomain is a suitable site for the incorporation of both AzF and BzF (23). Sequence alignment (Figure 2A) and structural superimposition (Figure 2B) of the ATAD2 and TRIM24 bromodomains revealed that N981 of ATAD2 and L922 of TRIM24 occupy analogous positions within the hydrophobic pocket. Based on this structural similarity, we selected L922 within the “LAF shelf” of TRIM24 as a site for the site-specific incorporation of AzF and BzF.

**Figure 2.**
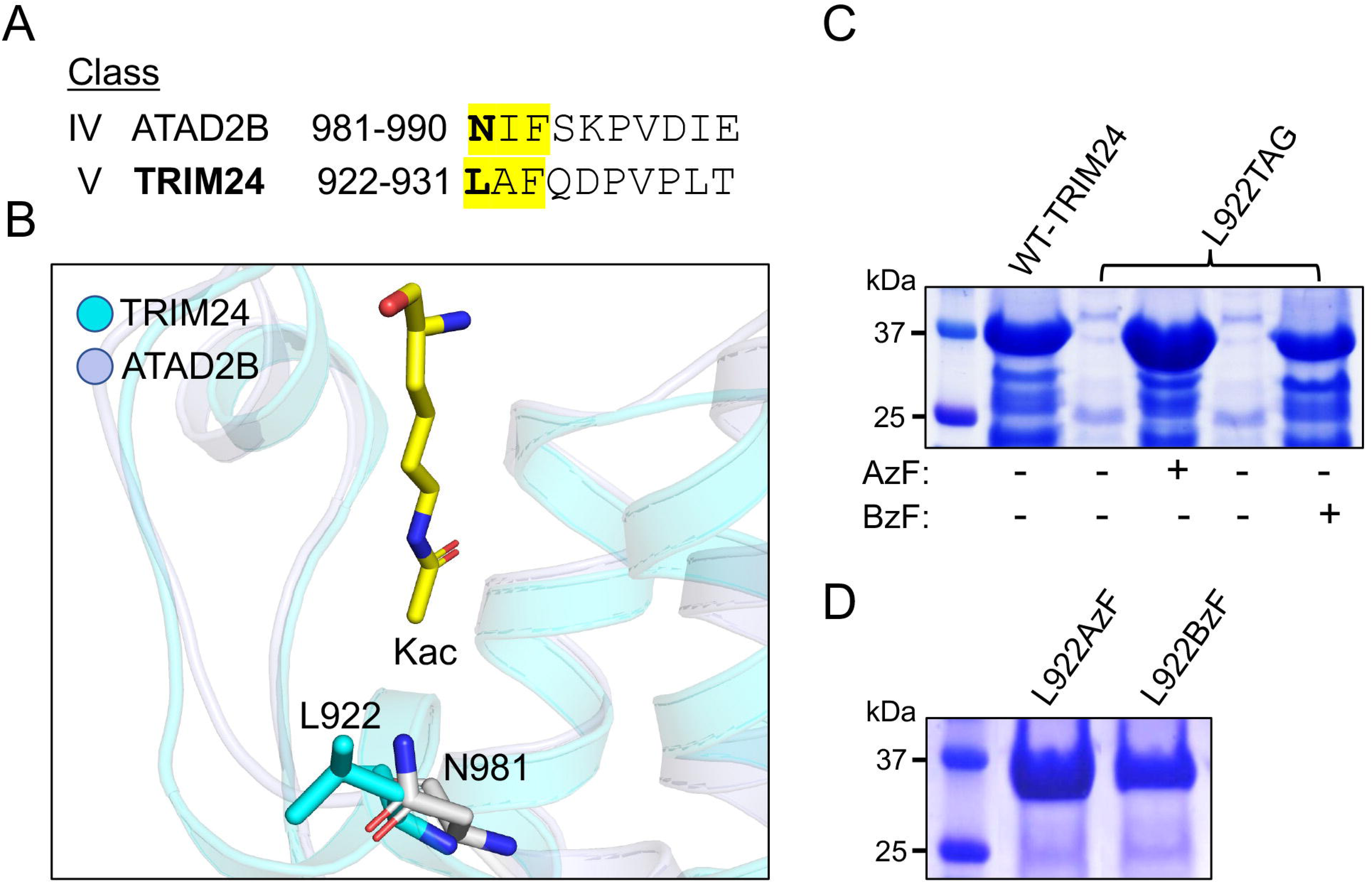
Engineering of the TRIM24 bromodomain with genetically encoded photocrosslinkable amino acids. (A) Alignment of partial bromodomain sequences of ATAD2B and TRIM24. The corresponding residues N981 in ATAD2B and L922 in TRIM24 are highlighted in yellow and shown in bold. (B) Structural superposition of the ATAD2B bromodomain (PDB ID: 3LXJ) and TRIM24 bromodomain (PDB ID: 3O34), showing the positions of N981 and L922 within the acetyl-lysine-binding pocket. L922 was selected as the site for incorporation of the photocrosslinkable amino acids 4-azido-L-phenylalanine (AzF) and 4-benzoyl-L-phenylalanine (BzF). (C) Coomassie blue-stained SDS–PAGE showing expression of wild-type TRIM24 bromodomain and the L922TAG mutant in Escherichia coli in the absence or presence of AzF or BzF. (D) Coomassie blue-stained SDS-PAGE showing the expression and purification of the engineered TRIM24 bromodomain variants L922AzF and L922BzF.

To enable site-specific incorporation of these unnatural amino acids, we introduced an amber stop codon (TAG) at position L922 of the TRIM24 PHD-Bromo module, generating the L922TAG mutant (Table S1 and Table S2). Amber suppression was achieved using the evolved orthogonal *Methanococcus jannaschii* TyrRS-tRNACUA^Tyr^ pair (25,26) (Table S1). The TRIM24 PHD-Bromo variants containing AzF or BzF at position 922 were expressed in *Escherichia coli* in the presence of the corresponding unnatural amino acid. Both variants were expressed at high levels and were subsequently purified by size-exclusion chromatography (Figure 2C,D and Figure S1).

We next evaluated whether incorporation of AzF or BzF at L922 affected the peptide-binding activity of the TRIM24 PHD-Bromo module. Isothermal titration calorimetry (ITC) was performed using the H3(1–33)K4K23ac peptide (Table S3). Wild-type TRIM24 PHD-Bromo bound the peptide with a *K*_D_ of 0.4 ± 0.05 μM, consistent with our previously reported affinity (17). The L922AzF variant retained binding activity, with a *K*_D_ of 1.0 ± 0.04 μM, whereas the L922BzF variant exhibited a modest reduction in affinity, with a *K*_D_ of 2.5 ± 0.02 μM (Figure 3). These results demonstrate that substitution of L922 with either AzF or BzF preserves the ability of the TRIM24 PHD-Bromo module to recognize the acetylated histone H3 peptide, supporting the use of these variants for subsequent photocrosslinking studies.

**Figure 3.**
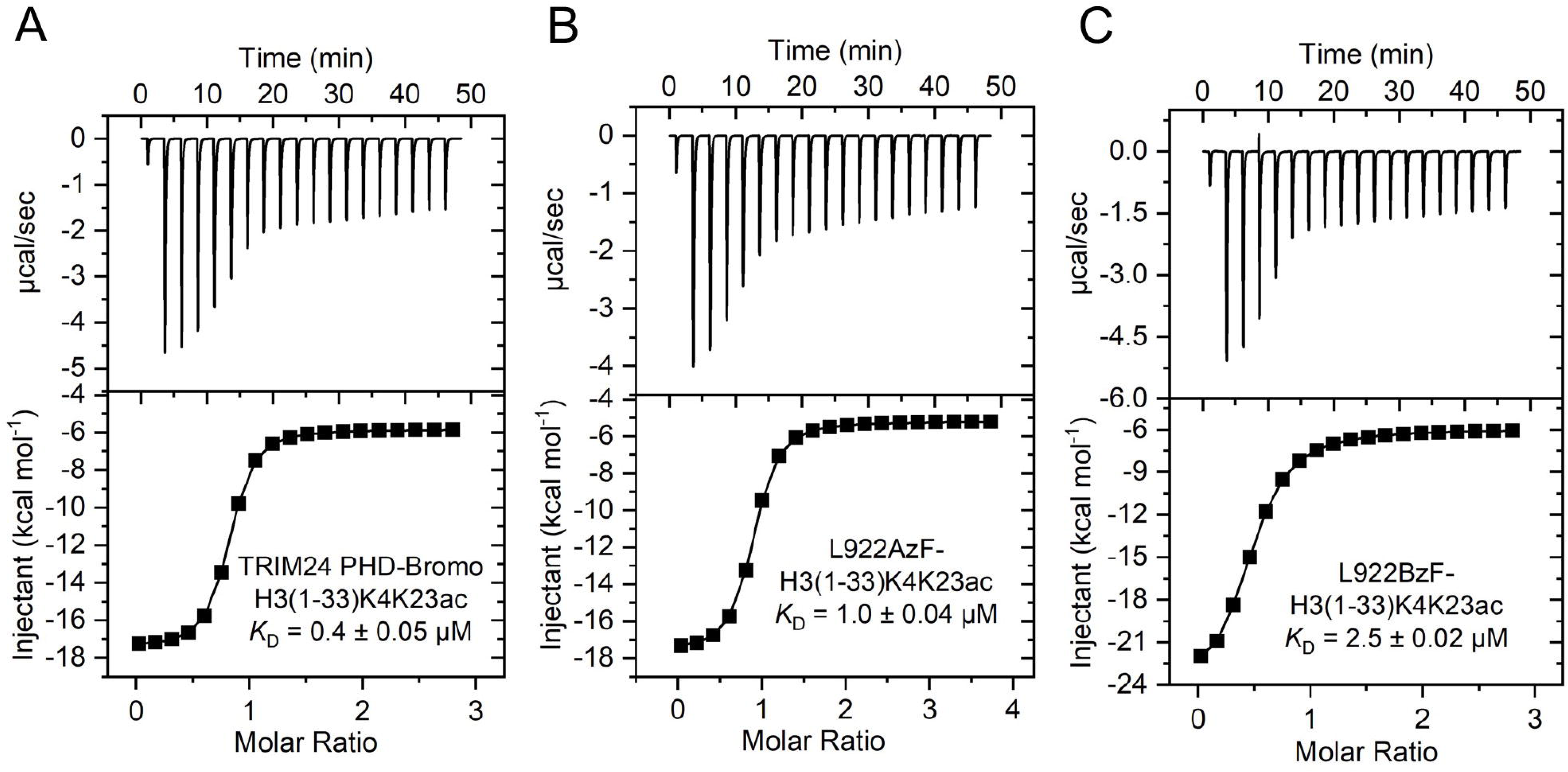
Isothermal titration calorimetry analysis of TRIM24 bromodomain binding to an acetylated histone H3 peptide. (A–C) Representative ITC profiles showing the binding of TRIM24 PHD–bromodomain (A) and the engineered bromodomain variants L922AzF (B) and L922BzF (C) to the H3(1–33)K4K23ac peptide. The upper panels show the raw heat-release profiles following successive peptide injections, and the lower panels show the corresponding integrated binding isotherms.

### Photocrosslinking of engineered TRIM24 bromodomain variants with acetylated histone H4 peptides

To evaluate the photocrosslinking capability of the engineered TRIM24 bromodomain variants L922AzF and L922BzF, we performed UV-induced photocrosslinking assays using acetylated histone H4 peptides. As illustrated in Figure 4A, the engineered TRIM24 bromodomain variants were incubated with TAMRA-labeled acetylated H4 peptides and irradiated with 365-nm UV light to induce covalent crosslinking between the photoreactive amino acid incorporated at L922 and the interacting peptide. Three acetylated H4 peptides, H4K12ac, H4K16ac, and H4Kac4, were used for these experiments. Each peptide was labeled with TAMRA at its C-terminus to enable fluorescence-based detection of the crosslinked products (Figure 4B).

**Figure 4.**
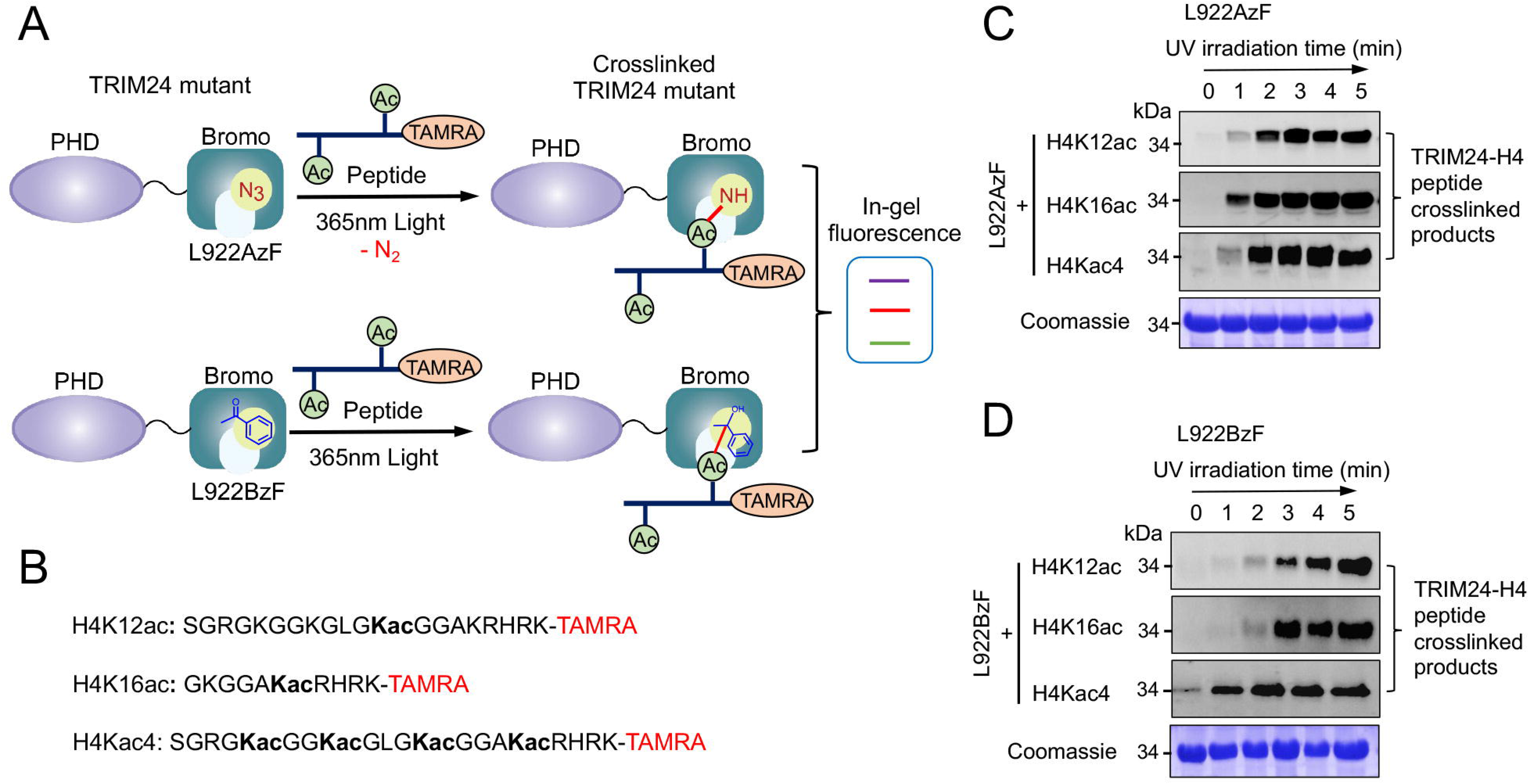
Photocrosslinking of engineered TRIM24 bromodomain with acetylated histone H4 peptides. (A) Schematic representation of photocrosslinking of the engineered TRIM24 bromodomain variants L922AzF and L922BzF with TAMRA-labelled acetylated H4 peptides upon irradiation with 365 nm UV light. (B) Amino acid sequences of the TAMRA-labelled H4 peptides H4K12ac, H4K16ac, and H4Kac4 used for photo-crosslinking experiments, with acetylated lysine residues indicated in bold and the TAMRA fluorophore shown in red. (C–D) Time-dependent photo-crosslinking of TRIM24-L922AzF (C) and TRIM24-L922BzF (D) with TAMRA-labelled H4 peptides following UV irradiation for the indicated times. Crosslinked products were detected by in-gel fluorescence, and Coomassie staining was used to assess protein loading.

We first examined the time-dependent photocrosslinking activity of the TRIM24-L922AzF variant with the three acetylated H4 peptides. Following UV irradiation, the reaction mixtures were subjected to Ni-NTA affinity enrichment, and the crosslinked products were detected by in-gel fluorescence. TRIM24-L922AzF exhibited clear UV-dependent crosslinking with all three acetylated H4 peptides, with the crosslinking signal increasing with prolonged irradiation (Figure 4C and Figure S2). Detectable crosslinked products were observed within approximately one min for H4K12ac, H4K16ac and H4Kac4, indicating that the L922AzF variant can undergo photo-induced covalent capture with multiple acetylated H4 peptides.

We next evaluated the photocrosslinking activity of the TRIM24-L922BzF variant under the same experimental conditions. Similar to L922AzF, L922BzF showed UV-dependent crosslinking with all three acetylated H4 peptides (Figure 4D and Figure S3). However, the time-dependent crosslinking profiles differed among the peptides. Crosslinking with H4K12ac and H4K16ac became detectable after approximately two min of UV irradiation, whereas H4Kac4 showed detectable crosslinking within approximately one min. The crosslinking signal increased with increasing irradiation time, demonstrating that the L922BzF variant is also capable of covalent capture of acetylated H4 peptides.

Taken together, these results demonstrate that site-specific incorporation of either AzF or BzF at L922 enables the TRIM24 bromodomain to undergo UV-induced covalent crosslinking with acetylated histone H4 peptides. Although both variants captured the H4 peptides, L922AzF generally showed an earlier onset of detectable crosslinking than L922BzF under the conditions tested. These findings establish L922AzF and L922BzF as suitable photocrosslinking variants for subsequent investigation of TRIM24 interactions with acetylated histone substrates and cellular proteins.

### Molecular modelling of TRIM24 L922AzF and L922BzF variants

The ITC and photocrosslinking experiments showed that incorporation of AzF or BzF at L922 preserves the ability of the TRIM24 bromodomain to interact with acetylated histone H4 peptides, although the two variants displayed distinct binding and crosslinking profiles. To investigate the structural basis for these differences, we performed molecular modelling and molecular dynamics (MD) simulations of the TRIM24 bromodomain variants in complex with the H4K16ac peptide. A crystal structure of the TRIM24 PHD-Bromo module in complex with the H4(14–19)K16ac peptide was obtained from the RCSB Protein Data Bank and used as the starting structure. The L922 residue was subsequently modelled as either AzF or BzF to generate the corresponding photo-reactive variants (Figure 5A).

**Figure 5.**
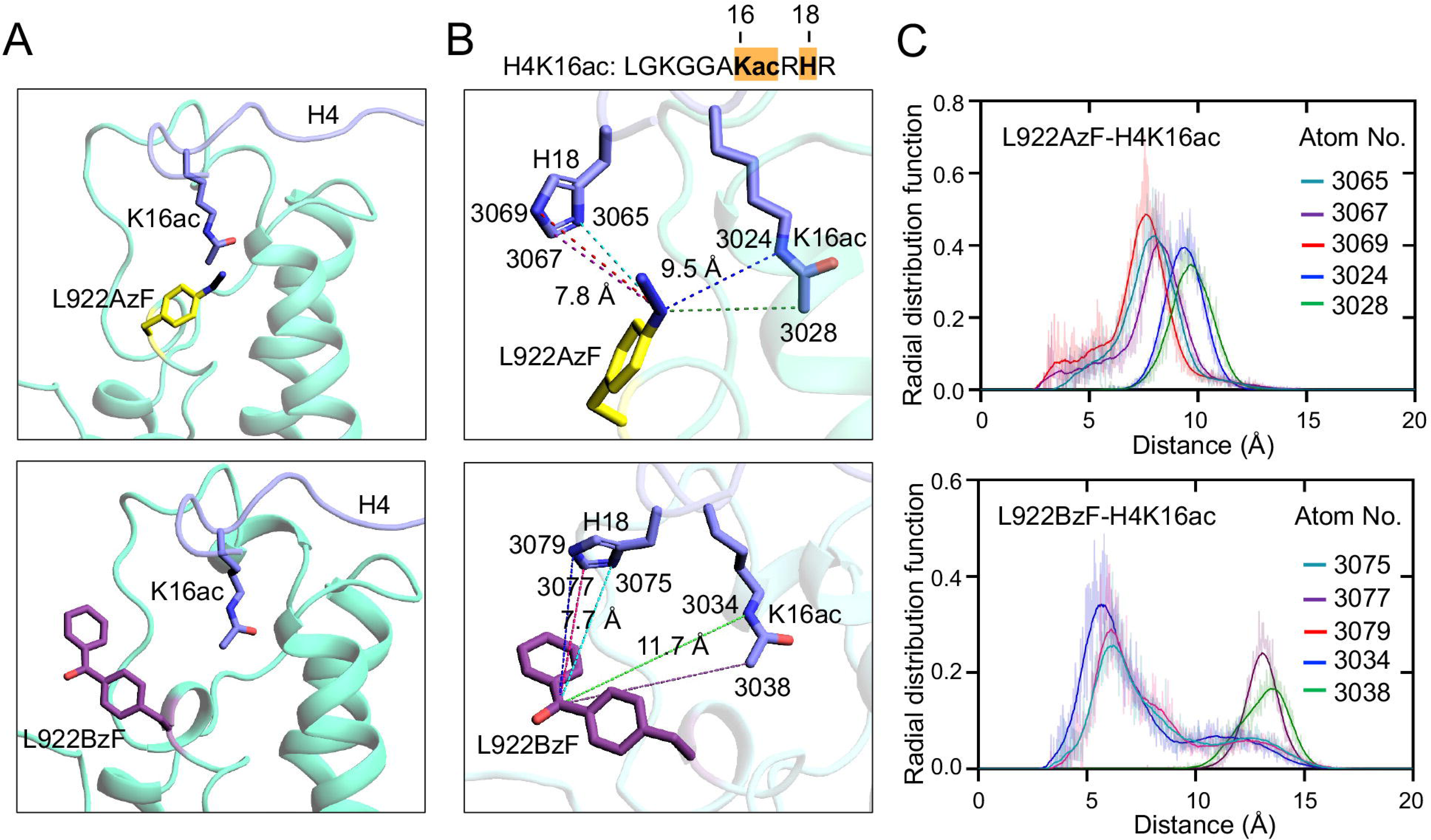
Molecular modelling of engineered TRIM24 bromodomain in complex with the H4K16ac peptide. (A) Structural models of the TRIM24 bromodomain containing L922AzF (yellow) or L922BzF (purple) in complex with the H4K16ac peptide. (B) Close-up views of the predicted photo-crosslinking sites showing the spatial proximity of the engineered residues to K16ac and H18 of the H4K16ac peptide. The indicated distances represent the distances between the corresponding atoms in the TRIM24 bromodomain and the H4K16ac peptide. (C) Radial distribution functions showing the distance-dependent spatial distribution of selected atoms in the L922AzF–H4K16ac and L922BzF–H4K16ac models. The atom numbers correspond to those indicated in the respective molecular models.

All-atom MD simulations were performed to examine the conformational behavior of the engineered residues and their spatial relationship with the H4K16ac peptide. Particular attention was given to the distances between the photoreactive groups at position 922 and atoms located in the vicinity of the acetylated lysine (K16ac) and H18 residues of the peptide (Figure 5B). Radial distribution function (RDF) analysis revealed distinct spatial distributions for the AzF- and BzF-containing variants (Figure 5C). For L922AzF, the azide-containing group showed a prominent distribution at approximately 7.8 Å from the H18-associated atoms, whereas the corresponding distance from atoms associated with H4K16ac was approximately 9.5 Å. These distances indicate that the AzF group can adopt conformations that place it in relatively close proximity to the peptide, particularly near H18.

In the L922BzF variant, the benzophenone moiety showed a closer spatial distribution with atoms associated with H18, with distances in the approximate range of 5–7 Å. In contrast, the distance between the benzophenone moiety and atoms associated with the acetylated K16 residue was substantially larger, reaching approximately 11.7 Å (Figure 5C). Thus, although the BzF group can approach H18 within the modelled complex, its greater separation from the acetylated lysine places the benzophenone moiety in a less favorable geometry for covalent capture of the K16ac region.

Together, the MD and RDF analyses provide a structural rationale for the distinct photo-crosslinking behavior observed for the two TRIM24 variants. The L922AzF group maintains relatively close proximity to the H4K16ac peptide across the simulated conformations, whereas the L922BzF group exhibits a more pronounced spatial separation from the acetylated lysine. These differences in the positioning of the photo-reactive groups may contribute to the distinct crosslinking profiles observed experimentally for L922AzF and L922BzF. Importantly, the modelling results support the use of L922AzF as a suitable probe for subsequent identification of TRIM24-interacting proteins.

### Photocrosslinking of engineered TRIM24 bromodomain mutants with the cellular proteome

Having established the photocrosslinking activity of the engineered TRIM24 bromodomain variants L922AzF and L922BzF with acetylated H4 peptides, we next investigated their ability to capture interacting proteins from the cellular proteome. HEK293T cells were treated with the deacetylase inhibitor trichostatin A to increase protein acetylation, and the resulting hyperacetylated cell lysates were incubated with purified TRIM24-L922AzF or TRIM24-L922BzF. Following incubation, the samples were irradiated with 365 nm UV light to induce covalent crosslinking. The His-tagged TRIM24 complexes were subsequently enriched by Ni-NTA affinity capture, followed by extensive washing to remove non-specifically associated proteins (Figures 6A and 7A).

**Figure 6.**
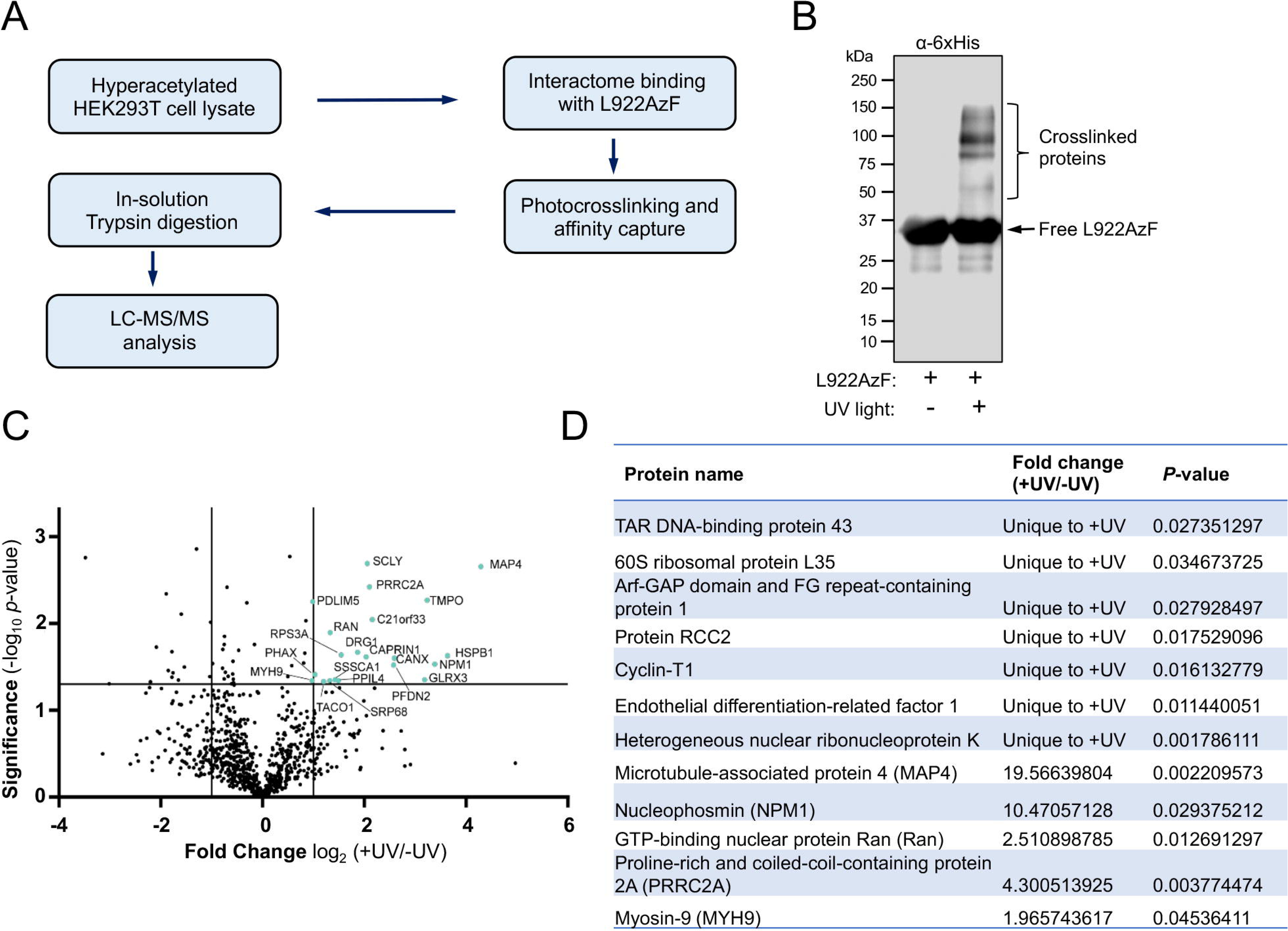
Photo-crosslinking of the TRIM24 bromodomain mutant L922AzF with the cellular proteome and identification of interacting proteins. HEK293T cells were treated with the deacetylase inhibitor trichostatin A (TSA) to generate a hyperacetylated proteome. (A) Schematic representation of the workflow for photo-crosslinking, affinity enrichment, and LC–MS/MS identification of proteins interacting with TRIM24-L922AzF. (B) Immunoblot analysis using an anti-6×His antibody showing UV-dependent formation of higher-molecular-weight crosslinked protein complexes with TRIM24-L922AzF. Free L922AzF and crosslinked proteins are indicated. (C) Volcano plot showing proteins enriched in the +UV sample relative to the −UV control. The x-axis represents log₂ fold change (+UV/−UV), and the y-axis represents statistical significance (−log₁₀ *P*-value); selected proteins are labeled. (D) Proteins identified as TRIM24-L922AzF-interacting proteins, with their fold changes and corresponding *P*-values.

**Figure 7.**
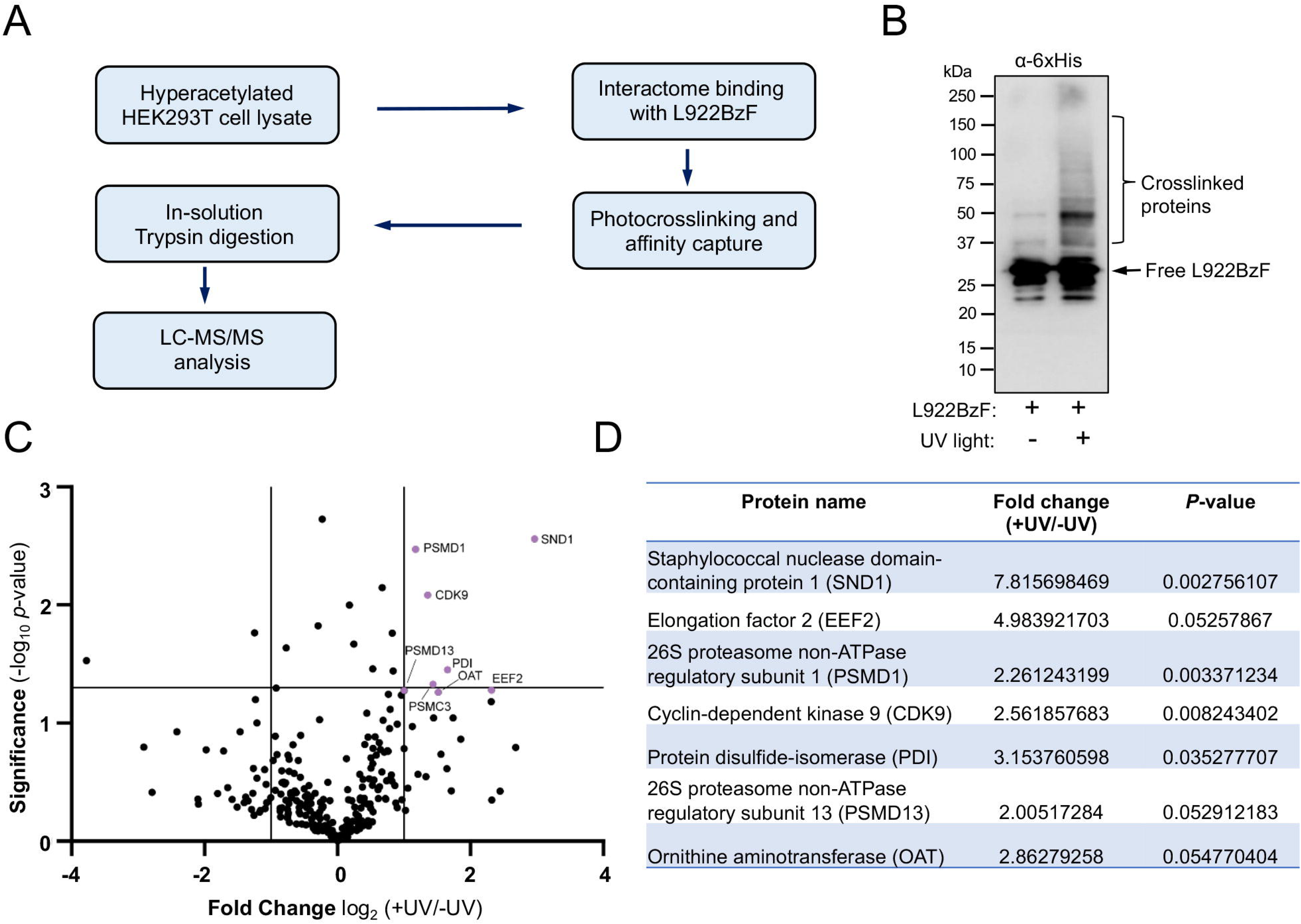
Photo-crosslinking of the TRIM24 bromodomain mutant L922BzF with the cellular proteome and identification of interacting proteins. HEK293T cells were treated with the deacetylase inhibitor trichostatin A (TSA) to generate a hyperacetylated proteome. (A) Schematic representation of the workflow for photo-crosslinking, affinity enrichment, and LC–MS/MS identification of proteins interacting with TRIM24-L922BzF. (B) Immunoblot analysis using an anti-6×His antibody showing UV-dependent formation of higher-molecular-weight crosslinked protein complexes with TRIM24-L922BzF. Free L922BzF and crosslinked proteins are indicated. (C) Volcano plot showing proteins enriched in the +UV sample relative to the −UV control. The x-axis represents log₂ fold change (+UV/−UV), and the y-axis represents statistical significance (−log₁₀ *P*-value); selected proteins are labeled. (D) Proteins identified as TRIM24-L922BzF-interacting proteins, with their fold changes and corresponding *P*-values.

For the L922AzF variant, UV irradiation resulted in the appearance of higher-molecular-mass species in addition to the band corresponding to free TRIM24-L922AzF, as detected by anti-6×His immunoblotting (Figure 6B). These higher-molecular-mass species were substantially reduced in the absence of UV irradiation, supporting UV-dependent covalent capture of proteins by the engineered bromodomain. A similar UV-dependent crosslinking pattern was observed with the L922BzF variant (Figure 7B), demonstrating that both photo-reactive amino acids enable covalent capture of proteins from the cellular proteome.

To identify proteins enriched by photocrosslinking, the affinity-purified material was subjected to in-solution trypsin digestion followed by LC-MS/MS analysis (Figures 6A and 7A). Two independent biological replicates were analyzed for both UV-irradiated and non-irradiated samples. Label-free quantification was performed using the MaxQuant LFQ algorithm (27) (IB_3), and the resulting datasets were processed to remove contaminants and quantify proteins enriched following UV irradiation. Relative intensity-based absolute quantification (riBAQ) values were calculated and averaged across biological replicates. Proteins showing increased abundance in the UV-irradiated samples were identified based on fold enrichment and statistical significance using a t-test, with a *P*-value threshold of ≤0.05.

For TRIM24-L922AzF, 75 proteins were detected exclusively in the UV-irradiated samples, whereas 70 proteins were detected exclusively in the non-irradiated samples, with 909 proteins detected under both conditions. Among the proteins common to both conditions, 21 showed significant enrichment following UV irradiation (Figure 6C,D). The enriched proteins included factors associated with transcription, RNA metabolism, nucleocytoplasmic transport, translation, cytoskeletal organization, and protein homeostasis. Notably, nucleophosmin (NPM1), and heterogeneous nuclear ribonucleoprotein K (HNRNPK) were among the proteins enriched following UV irradiation.

For TRIM24-L922BzF, 47 proteins were detected exclusively in the UV-irradiated samples and 18 proteins exclusively in the non-irradiated samples, while 284 proteins were detected under both conditions. Of the proteins common to the two conditions, eight showed significant enrichment following UV irradiation (Figure 7C,D). These included proteins associated with transcription, RNA processing, translation, cytoskeletal organization, and protein homeostasis. Several proteins, including NPM1 and CDK9, were identified in the datasets obtained using both L922AzF and L922BzF, suggesting that these proteins may represent reproducible candidate interactors of the TRIM24 bromodomain.

Together, these results demonstrate that site-specific incorporation of AzF or BzF at L922 enables UV-dependent covalent capture of proteins from the cellular proteome. The identification of both shared and variant-specific proteins provides a set of candidate TRIM24-associated proteins for further investigation. In particular, the enrichment of proteins involved in transcription and RNA-associated processes is consistent with a broader role for TRIM24 in cellular regulatory networks beyond its established chromatin-associated functions.

### CDK9, NPM1 and TRIM24 colocalize at transcription start sites of cancer-associated hub genes

The proteomic analysis identified a broad range of proteins associated with the TRIM24 bromodomain. To investigate whether selected interactors share genomic occupancy with TRIM24, we performed ChIP-seq analysis of NPM1 and CDK9. NPM1 (GSM6775431) was selected from the TRIM24-L922AzF interactome, whereas CDK9 (GSM2405924) was selected from the TRIM24-L922BzF interactome. The corresponding ChIP-seq datasets were retrieved from the Gene Expression Omnibus (GEO) database and processed using the Galaxy platform. Following adapter trimming with Trimmomatic, the reads were aligned to the reference genome using Bowtie2, and enriched genomic regions were identified using MACS2 peak calling. ChIP-seq data for TRIM24 (ENCSR957LDM) were obtained from the ENCODE database. The processed datasets were subsequently visualized using the Integrative Genomics Viewer (IGV) to examine the genomic distribution and co-localization of TRIM24, CDK9, and NPM1.

We first examined cancer-associated hub genes identified in breast cancer and observed that TRIM24, CDK9, and NPM1 showed overlapping enrichment at or near the transcription start sites of ANLN, PLK1, CDK1, and NME1 (Figure 8A). Similar co-localization was observed at the transcription start sites of prostate cancer-associated hub genes, including *CCNB2*, *CDK1*, *PLK1*, and *CDKN3* (Figure 8B). In colorectal cancer-associated hub genes, *TRIM24*, *CDK9*, and *NPM1* likewise exhibited overlapping occupancy at the transcription start sites of HSPD1, TRA2A, RBM39, and CDK4 (Figure 8C). These observations indicate that TRIM24, CDK9, and NPM1 show overlapping occupancy at the transcription start sites of multiple cancer-associated hub genes, supporting a potential role for these proteins in transcriptional regulation.

**Figure 8.**
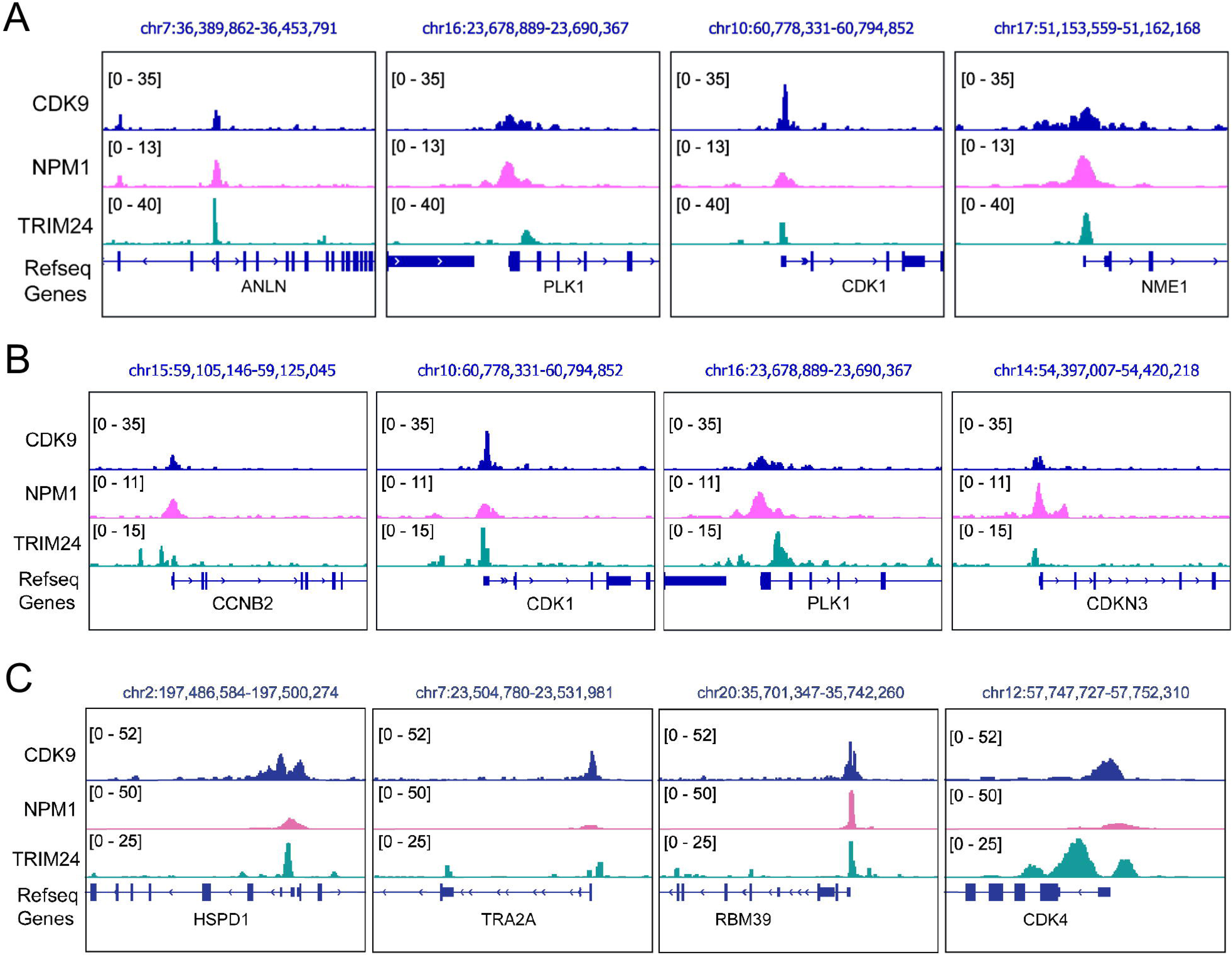
Co-localization of CDK9, NPM1, and TRIM24 at transcription start sites of cancer-associated hub genes. IGV snapshots showing the genomic binding profiles of CDK9, NPM1, and TRIM24 at or near the transcription start sites of selected hub genes associated with (A) breast cancer (ANLN, PLK1, CDK1, and NME1), (B) prostate cancer (CCNB2, CDK1, PLK1, and CDKN3), and (C) colorectal cancer (HSPD1, TRA2A, RBM39, and CDK4). RefSeq gene annotations are shown below the corresponding genomic tracks.

To further investigate the functional relevance of these proteins, we performed KEGG pathway enrichment analysis using the ClusterProfiler package (28)(IB_8) within the Bioconductor framework. The analysis identified enrichment in several pathways, including thyroid hormone signalling, Rap1 signalling, FoxO signalling, MAPK signalling, and Hippo signalling (Figure S5). Together, these findings suggest that the TRIM24-associated proteins CDK9 and NPM1 may participate in transcriptional regulatory networks linked to multiple cancer-relevant cellular pathways.

## Conclusion

In this study, we established a site-specific photocrosslinking strategy to investigate the interaction landscape of the TRIM24 bromodomain. Photocrosslinkable amino acids, AzF and BzF, were successfully incorporated at L922 within the acetyllysine-binding pocket of TRIM24, and both engineered variants retained the ability to recognize an acetylated histone H3 peptide. Photocrosslinking experiments further demonstrated that both L922AzF and L922BzF underwent UV-dependent covalent capture of multiple acetylated histone H4 peptides, although L922AzF generally exhibited an earlier onset of detectable crosslinking. Molecular modelling and molecular dynamics simulations revealed distinct spatial distributions of the AzF and BzF groups within the TRIM24-H4K16ac complex, providing a structural rationale for the different crosslinking profiles observed experimentally.

We subsequently applied the L922AzF and L922BzF variants to photocrosslinking-based proteomic profiling of HEK293T cellular extracts. Quantitative LC-MS/MS analysis revealed distinct protein capture profiles for the two probes and identified a diverse set of candidate nonhistone proteins associated with the TRIM24 bromodomain. NPM1 was enriched in the L922AzF dataset, whereas CDK9 was enriched in the L922BzF dataset, together with additional candidate proteins including RAN, EEF2, PRRC2, MYH9, MAP4, and PSMD1. ChIP-seq analysis further revealed overlapping genomic occupancy of TRIM24, CDK9, and NPM1 at the transcription start sites of hub genes associated with breast, prostate, and colorectal cancers. These findings provide a genomic context in which the identified candidate proteins may be associated with TRIM24-dependent transcriptional regulation.

Collectively, these findings demonstrate that site-specific incorporation of photocrosslinkable amino acids at the TRIM24 bromodomain can preserve acetyllysine recognition while enabling covalent capture of acetylated histone peptides and candidate nonhistone proteins. The distinct proteomic profiles obtained with AzF and BzF highlight the potential of complementary photocrosslinking chemistries to probe diverse protein association landscapes. More broadly, coupling site-specific photocrosslinking with quantitative proteomics provides a complementary approach for extending the study of epigenetic reader domains beyond canonical histone recognition and accessing protein associations that may be difficult to capture using conventional enrichment approaches. Although the candidate protein associations and their functional relationships with TRIM24 require further biochemical and cellular validation, this study provides a framework for defining the nonhistone interaction landscape of TRIM24 and investigating its potential contributions to transcriptional regulation and cancer-relevant cellular processes.

## Materials and methods

### General materials, methods, and equipment

All of the plasmids for bacterial expression were obtained as gifts from individual laboratories or purchased from Addgene (Table S1). Mutagenic primers were obtained from Sigma-Aldrich (Table S2). Commercially available competent bacterial cells were used for protein expression and mutagenesis. HEK293T cells were obtained from the American Type Culture Collection and used according to the manufacturer’s protocol. All of the antibodies used in this study were purchased from established vendors and used following the manufacturer’s protocol. AzF was a kind gift from K. Islam (University of Pittsburgh), synthesized and characterized as reported previously (20). BzF was purchased from BLD Pharmatech (catalog no. BD135722). All histone peptides were synthesized by GL Biochem Ltd and S. Biochem Ltd and purified by HPLC to 98% purity (Table S3 and Figures S6-S13). The concentration of unlabeled peptide was determined on the basis of the observation that 1 mg/ml peptide generates an absorbance (A_205_) of 30 at 205 nm. The concentrations of TAMRA-labelled peptides were determined by measuring the absorbance at a wavelength of 555 nm with an extinction coefficient of 65,000 L^-1^ cm^-1^ M^-1^. MALDI-MS confirmed the integrity of the purified peptides.

### Mutagenesis, expression, and purification of the TRIM24 PHD-bromodomain

Protein expression and purification were performed using previously described methods (11,17). The N-terminal 6×His-TRIM24 PHD-bromodomain plasmid (Addgene, catalog no. 74647) was transformed into One Shot BL21 Star (DE3) *Escherichia coli* competent cells (Invitrogen, catalog no. C601003) using pET15b ampicillin-resistant vectors. A single colony was picked and grown overnight at 37 °C in 10 ml of LB broth in the presence of 50 μg mL^-1^ ampicillin. The culture was diluted 100-fold and allowed to grow at 37 °C to an absorbance (A600) of 1. Protein expression was induced overnight at 17 °C with 0.5 mM IPTG in an Innova 44 Incubator shaker (New Brunswick Scientific). Proteins were purified as follows. Harvested cells were resuspended in 15 ml of lysis buffer (50 mM HEPES [pH 7.5], 300 mM NaCl, 5 mM β-mercaptoethanol, 5% glycerol, 25 mM imidazole, lysozyme, DNase, and 1:200 (v/v) Protease Inhibitor Cocktail III [Calbiochem]). The cells were lysed by pulse sonication and centrifuged at 13,000 rpm for 40 min at 4 °C. According to the manufacturer’s instructions, the soluble extracts were subjected to Ni-NTA agarose resin (QIAGEN, catalog no. 30210). After 20 volumes of wash buffer (50 mM HEPES pH 7.5, 300 mM NaCl, 5 mM β-mercaptoethanol, 5% glycerol, and 25 mM imidazole) had been passed through the column, proteins were eluted with a buffer containing 50 mM HEPES pH 7.5, 300 mM NaCl, 5 mM β-mercaptoethanol, 5% glycerol, and 250 mM imidazole. Proteins were further purified by gel-filtration chromatography (Superdex-75) using an AKTA pure FPLC system (GE Healthcare) with a buffer containing 50 mM HEPES pH 7.5, 200 mM NaCl, and 5% glycerol. Purified proteins were concentrated using an Amicon Ultra-10k centrifugal filter device (Merck Millipore), and the concentration was determined using a Bradford assay kit (Bio-Rad Laboratories) with bovine serum albumin as a standard. The proteins were aliquoted and stored at −80 °C until use.

Site-directed mutagenesis was performed to generate the TRIM24 bromodomain amber codon (TAG) mutant L922TAG using the QuikChange Lightning Site-Directed Mutagenesis Kit, and the resulting mutant plasmids were confirmed by DNA sequencing. To express TRIM24 bromodomain mutants carrying AzF or BzF at the L922 position, the TRIM24 amber variant (L922TAG) was co-transformed with pEVOL-pAzF (Addgene, catalog no. 31186) (26) or pEVOL-pBzF (Addgene, catalog no. 31190) (25) into One Shot BL21 Star (DE3) *E. coli* competent cells. The pEVOL plasmids were kind gift from K. Islam, University of Pittsburgh. After transformation, cells were recovered in 200 μl of SOC medium and incubated in a 37 °C shaker for 1 h at 225 rpm before being plated on a dual antibiotic LB Miller agar plate containing 100 μg mL^-1^ ampicillin and 35 μg mL^-1^ chloramphenicol. A single colony was picked and inoculated into 5 mL of LB Miller broth in the presence of 100 μg mL^-1^ ampicillin and 35 μg mL^-1^ chloramphenicol and grown overnight in a 37°C shaker at 225 rpm. This overnight culture was centrifuged (Eppendorf Centrifuge 5910 R, S-4xUniversal swing-bucket rotor) at 1000 × g (2158 rpm) for 10 min at room temperature (RT), and the supernatant (LB Miller broth) was discarded. The collected cell pellet was then resuspended in 1 mL of M9 medium and used to inoculate 500 mL of GMML medium supplemented with 100 μg mL^-1^ ampicillin and 35 μg mL^-1^ chloramphenicol. Cells were allowed to grow at 37 °C in an incubator shaker (225 rpm) to an optical density (OD600) of 1. The unnatural amino acid AzF was prepared by dilution in 5mL of sterilized deionized water, and BzF was dissolved in 1mL of sterilized 1 M NaOH; the solutions were then added, respectively, to the bacterial culture to a final concentration of 1 mM. The cells were then cooled to 17 °C and allowed to grow for 30 min; at this stage, expression of the AzF/BzF-specific aminoacyl-tRNA synthetase was induced with 0.05% (w/v) arabinose, and the cells were incubated with shaking for an additional 30 min at 17 °C. Finally, TRIM24 protein expression was induced by the addition of 0.5 mM IPTG, and the cells were allowed to grow for 20 h at 17 °C. The TRIM24 bromodomain mutants were purified as described above while minimizing exposure to ambient light.

### Isothermal titration calorimetry

ITC binding experiments were performed as previously described (17). Briefly, experiments were carried out on a MicroCal PEAQ ITC instrument (Malvern) at 15 °C while the sample was being stirred at 750 rpm. Proteins and peptide samples were prepared in matching buffer containing 50 mM HEPES (pH 7.5), 200 mM NaCl, and 5% glycerol. Each titration comprised one initial injection of 0.4 μL lasting 0.8 s, followed by 18 injections of 2 μl lasting 4 s each at 2.5 min intervals. The initial injection was discarded during data analysis. The microsyringe (40 μL) was loaded with a peptide sample solution at a concentration of 1.5 to 1.8 mM. The peptide solution was injected into the 200 μL cell protein at a concentration of 75-100 μM. All of the data were fitted to a single-binding site model using the MicroCal PEAQ ITC analysis software to calculate the stoichiometry (*N*), binding constant (*K*_D_), enthalpy (*ΔH*), and entropy (*ΔS*) of the interaction. The final titration figures were prepared using OriginPro 2020 (OriginLab).

### Photocrosslinking experiment with histone peptides and in-gel fluorescence

Photocrosslinking of TRIM24 bromodomain mutants (L922AzF or L922BzF) with acetylated histone peptides was performed as previously described (23). Briefly, 5 μM of H4 peptides (H4K12ac, H4K16ac, and H4Kac4) labelled with tetramethylrhodamine (TAMRA) at the C-terminus were pre-incubated with 50 μM TRIM24 bromodomain variants in 100 μL binding buffer (50 mM HEPES pH 7.5, 150 mM NaCl, and 0.001% Tween 20) for 30 min at 4 °C on a rotator. After incubation, the samples were split into two 50 μL volumes in clear PCR tubes (Axygen). The PCR tubes containing the sample were placed into semi-micro visible cuvettes (300 nm – 900 nm, Eppendorf, catalog no. 0030079353) filled with ice cubes to maintain the temperature and irradiated with ultraviolet light (365 nm, 8 W lamps, Model: ECX-F20.L V1, Vilber Lourmat, France) for different time intervals (0-5 min) at 4 °C. Negative-control samples were not subjected to UV irradiation and were stored in the dark at 4 °C. After irradiation, the samples were transferred into 1.5 mL tubes containing 50-60 μL of Ni-NTA agarose beads (QIAGEN) pre-equilibrated with the buffer (50 mM HEPES pH 7.5, 150 mM NaCl, and 0.001% Tween-20), and the samples were incubated for 1 h at 4 °C on a rotator. The uncrosslinked histone peptides were removed by washing the samples with wash buffer (50 mM HEPES pH 7.5, 400 mM KCl, 1% Triton X-100) five times at room temperature. The proteins were eluted with 30 μL elution buffer containing 50 mM HEPES pH 7.5 and, 500 mM imidazole and incubated for 5 min at room temperature with intermittent mixing. The eluted proteins were mixed with 10 μL of 1 × TruPAGE LDS sample buffer (Sigma catalog no. PCG3009) and heat-denatured at 100 °C for 10 min. Finally, the crosslinked proteins were separated on a 15% SDS-PAGE gel and imaged on a G: BOX Chemi XRQ gel doc system (Syngene) at the TAMRA fluorophore excitation wavelength. The gel was subsequently stained with Coomassie brilliant blue R-250 staining solution to confirm the presence of proteins in all the samples.

### Molecular modelling and molecular dynamics simulations

Briefly, the crystal structure of the TRIM24 PHD-Bromo complex with the H4(14–19)K16ac peptide was obtained from the RCSB Protein Data Bank (PDB entry 3O36). The L922 residue of the TRIM24 bromodomain was individually replaced with AzF using the PyMOL Builder and with BzF using the SwissSidechain plugin in PyMOL. The resulting L922AzF and L922BzF variants were subjected to all-atom MD simulations using GROMACS 2018.3 (29) with the CHARMM27 all-atom force field (30), modified according to parameters developed by Smith et al. (31). The protein-peptide complexes were solvated in a dodecahedral box with a minimum distance of 1 nm between the protein and the box boundary. Sodium ions were added to neutralize the systems. Energy minimization was performed using the steepest-descent algorithm for 5000 steps, with a convergence criterion of <1000 kJ mol^-1^ nm^-1^. LINCS constraints were applied to constrained bonds during the simulations. Following energy minimization, the systems were equilibrated under NVT and NPT ensembles and subsequently subjected to 100-ns production MD simulations following the same steps described above. Radial distribution functions (RDFs) were calculated from the MD trajectories to examine the spatial distribution of the photo-reactive groups following AzF or BzF incorporation at L922. Distances between the azide moiety of AzF or the benzophenone moiety of BzF and the nearest atoms of the H4K16ac peptide were calculated. MD snapshots were extracted and analyzed using PyMOL. All plots were generated using GraphPad Prism.

### Mammalian cell culture

HEK293T cells were grown in Dulbecco’s modified Eagle’s medium supplemented with 10% FBS and antibiotics (penicillin/streptomycin cocktail) in a humidified atmosphere containing 5% CO2. Cells at 90% confluence were treated with 2 μM of the histone deacetylase inhibitor, trichostatin A (Sigma, catalog no. T8552) dissolved in dimethyl sulfoxide to generate hyper-acetylated histones. Twenty hours post-treatment, the medium was removed, and the cells were rinsed with cold PBS. The harvested cells were rewashed with cold PBS and frozen as a dry pellet at −80 °C.

### Preparation of HEK293T cell lysate

Twenty hours post-treatment with the deacetylase inhibitor (Trichostatin A), the medium was removed and the HEK293T cells were rinsed with cold PBS buffer. The harvested cells were resuspended in ice-cold lysis buffer containing 25 mM Tris–HCl pH 7.4, 150 mM NaCl, 1% NP-40, 1 mM EDTA, 5% glycerol (Pierce IP lysis buffer, catalog no. 87787), and supplemented with 1× Pierce protease inhibitor cocktail; the cells were then incubated on ice for 5 min with periodic mixing. The cell lysate was centrifuged at 13,000 × g for 10 min at 4 °C to pellet the cell debris. The supernatant was transferred into a fresh microcentrifuge tube, and the protein concentration was determined by Bradford assay (Bio-Rad Laboratories). The supernatant containing cell lysate was used for subsequent photo-crosslinking and Western blotting experiments.

### Photocrosslinking with HEK293T cell lysate and western blotting

For photocrosslinking studies, 1 mg of HEK293T cell lysate was incubated with 50 μM of TRIM24-L922AzF or TRIM24-L922BzF in a buffer containing 50 mM HEPES pH 7.5, 150 mM NaCl, and 0.001% Tween 20. After 1 h of incubation at RT, the samples were subjected to UV irradiation at 365 nm for 30 min at 4 °C. Negative controls were not subjected to UV exposure. Samples were then bound to Ni-NTA agarose resin and incubated for 1 h at 4 °C with gentle rotation. To remove uncrosslinked proteins present in cell lysates, samples were washed five times with washing buffer (50 mM HEPES pH 7.5, 400 mM KCl, 5% Triton X-100). The crosslinked proteins were eluted in 30 μl elution buffer containing 50 mM HEPES pH 7.5 and 500 mM imidazole and incubated for 5 min at RT with intermittent mixing. The eluted proteins were separated on a 4 to 15% SDS-PAGE gel (Bio-Rad, catalog no. 4561085) and transferred onto a 0.45 μm PVDF membrane at a constant voltage of 80 V for 1.5 h at 4 °C. The membrane was rinsed in TBST buffer (50 mM Tris pH 7.4, 150 mM NaCl, and 0.1% Tween-20) and blocked for an hour at RT in 5% milk buffer prepared in TBST. Immunoblotting was done with anti-6×His tag antibody (Invitrogen, catalog no. MA121315) overnight at 4 °C. The membrane was washed with TBST buffer thrice at RT for 5 min each. The blots were then incubated with the horseradish peroxidase-conjugated secondary antibody, goat anti-mouse IgG (Invitrogen, catalog no. 31431) at a 1:5000 dilution in TBST containing 5% non-fat dry milk. The membrane was rewashed with TBST buffer thrice at RT for 5 min each. Protein bands were visualized by chemiluminescence using SuperSignal West Pico PLUS substrate (Invitrogen, catalog no. 34577) following the manufacturer’s protocol.

### LC-MS/MS analysis

In-solution trypsin digestion was carried out according to the standard protocol. Proteolytic peptides from in-solution trypsin digestion were analyzed by nanoflow reversed-phase LC-MS/MS. Tryptic peptides were loaded onto a PepMap RSLC C18 column (2 μm, 100 Å × 50 cm) of the LC system (Q-Exactive Plus Biopharma, Thermo Fisher Scientific). Chromatographic separation was performed using a binary solvent system (solvent A: water with 0.1% formic acid; solvent B: acetonitrile and water at a ratio of 85:15 with 0.1% formic acid). MaxQuant v2.0.3.0 software (32) (https://www.maxquant.org) was used for protein identification and quantitation. Mass spectrometry data were searched against the UniProt-proteome-UP000005640 using Andromeda. Protein identifications were reported with a false discovery rate of 1%. We used MaxQuant to calculate iBAQ, a measure of protein abundance.

The proteomics pipeline described by Krey et. al. was used with slight moderation for further data analysis (33). The “proteinGroups.txt” file obtained from MaxQuant was processed using Mathematica (v10) (https://www.wolfram.com/mathematica/). The software was used to (1) customize protein nomenclature by replacing default names and symbols with user-specified designations; (2) filter out entries designated by MaxQuant as “Reverse” or “potential contaminant” while retaining proteins designated as “only identified by site”; (3) cluster proteins into groups in which more than 30% of their peptide sequences overlapped; and (4) generate an output file. The output file was imported, and the iBAQ value of each protein was divided by the total iBAQ value of the respective group to calculate the riBAQ value.

### Obtaining public ChIP-seq datasets

We looked into ENCODE server in order to obtain the processed TRIM24 ChIP-seq data for K562 cells (accession no. ENCSR957LDM). The unprocessed ChIP-seq data files for NPM1 (GSM6775431) in OCI-AML3 cells and CDK9 (GSM2405924) in MV4 cells were downloaded from Gene Expression Omnibus (GEO).

### ChIP-seq data processing and analysis

Raw ChIP-seq FASTQ files were retrieved and processed using the Galaxy platform (34) and in-house bioinformatics tools (35). Sequencing quality was assessed using FastQC, and adapter sequences were removed using Trimmomatic (36). The trimmed paired-end reads were aligned to the human reference genome (GRCh38) using Bowtie2 v2.4.2 (37). Peaks were identified from the aligned reads using MACS2 v2.1.1, with the corresponding input sample used as the background control (38). Peak reproducibility was assessed using irreproducible discovery rate (IDR) analysis (39), and peaks with an IDR ≤0.05 were retained as high-confidence peaks. Peaks overlapping ENCODE-defined blacklisted genomic regions (40) were excluded from further analysis. The processed ChIP-seq datasets were visualized using the Integrative Genomics Viewer (IGV) (41) to examine genomic binding profiles and co-localization. Downstream analyses were performed using the R/Bioconductor framework (v3.4.0), with the ChIPseeker package (42) used for genomic annotation and visualization of ChIP-seq peaks (43).

### Supporting Information

The Supporting Information includes plasmid construct information, mutagenesis primers, and a list of histone peptide sequences (Tables S1–S3), as well as Figures S1–S13. The figures include SDS-PAGE gels, photocrosslinking data, ITC binding data, peptide characterization by ESI-MS, and HPLC purity traces of peptides.

## Data Availability

The authors declare that the main data supporting the findings of this study are available within the article and its Supplementary materials.

## Author Contributions

B.S. conceived the ideas and supervised this study. I.B., and B.S. designed the experiments. I.B., S.C., and N.S.P. expressed and purified proteins and performed biochemical experiments. I.B. and B.S. analyzed data. B.S. wrote the paper.

## Acknowledgments

The authors acknowledge the infrastructural facilities supported by Central Instrumentation Facility (CIF), IISER Kolkata, and DST-FIST (SR/FST/LS-II/2017/93). I.B. acknowledges, CSIR-UGC, Government of India, for her doctoral fellowship.

## Abbreviations

PTM: posttranslational modification
TRIM24: tripartite motif-containing 24
PHD: plant homeodomain
BD: bromodomain
ITC: isothermal titration calorimetry
AzF: 4-azido-L-phenylalanine
BzF: 4-benzoyl-L-phenylalanine
IBPP: interaction-based protein profiling

