## Supplemental Figures for "Mapping the Nonhistone Interactome of the TRIM24 Bromodomain Using Genetically Encoded Complementary Photocrosslinkers"

**Table of Contents**

1. Supplementary Tables S2
2. Supplementary Figures S3-S11

**Table S1.** List of the genes used in the current study. The expression vector, antibiotic resistance and the affinity tag present for protein purification are provided.

| Gene Name | Vector | Affinity tag | Resistance |
|---|---|---|---|
| TRIM24 PHD-Bromodomain | pET15b | N-6xHis | Ampicillin |
| *M. j.* TyrRS-tRNA$_{CUA}^{Tyr}$ pair | pEVOL-pAzF | None | Chloramphenicol |
| *M. j.* TyrRS-tRNA$_{CUA}^{Tyr}$ pair | pEVOL-pBpF | None | Chloramphenicol |

**Table S2.** The sequence of the forward primer designed for site-directed mutagenesis. Reverse primers used are the reverse-complement to the given forward primer.

| TRIM24 Bromodomain mutant | Primer sequence |
|---|---|
| L922TAG | CTGCCATGAAATGAGCTAGGCTTTTCAAGACCCTG |

**Table S3.** List of histone peptides varying acetylation sites were synthesized and used for this study.

| Peptide | Peptide Sequence | Molecular weight (Da) |
|---|---|---|
| H3(1-33)K23ac | ARTKQTARKSTGGKAPRKQLAT**Kac**AARKSAPATG | 3436.92 |
| H4(1-20)K12ac | SGRGKGGKGLGKacGGAKRHRK**-TAMRA** | 2171.47 |
| H4(11-20)K16ac | GKGGA**Kac**RHRK**-TAMRA** | 1548.88 |
| H4(1-20)Kac4 (K5/K8/K12/K16) | SGRG**Kac**GG**Kac**GLG**Kac**GGA**Kac**RHRK**-TAMRA** | 2572.89 |

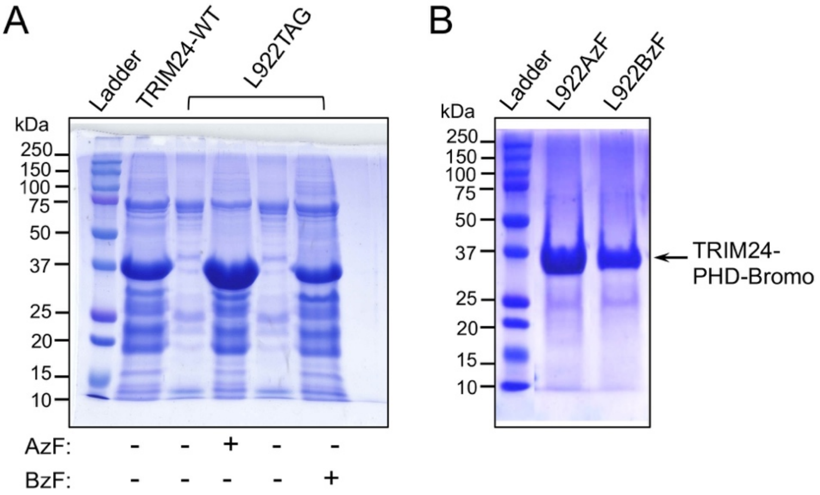


**Figure S1.** Figure S1. Engineering the TRIM24 PHD-Bromo module for site-specific incorporation of photocrosslinkable unnatural amino acids. (A) Coomassie-stained SDS–PAGE gel showing expression of the wild-type TRIM24 PHD-Bromo module and the L922TAG mutant in the absence (−) and presence (+) of 4-azido-L-phenylalanine (AzF) or 4-benzoyl-L-phenylalanine (BzF). A prominent band at ~33–35 kDa corresponds to the full-length TRIM24 PHD-Bromo module. The L922TAG mutant shows efficient full-length protein expression only in the presence of AzF or BzF, demonstrating successful amber suppression and incorporation of the corresponding unnatural amino acid. (B) Coomassie-stained SDS–PAGE gel of purified TRIM24 PHD-Bromo module containing AzF or BzF at position 922 (L922AzF and L922BzF, respectively). Both purified variants show a prominent band at the expected molecular weight of ~33–35 kDa.

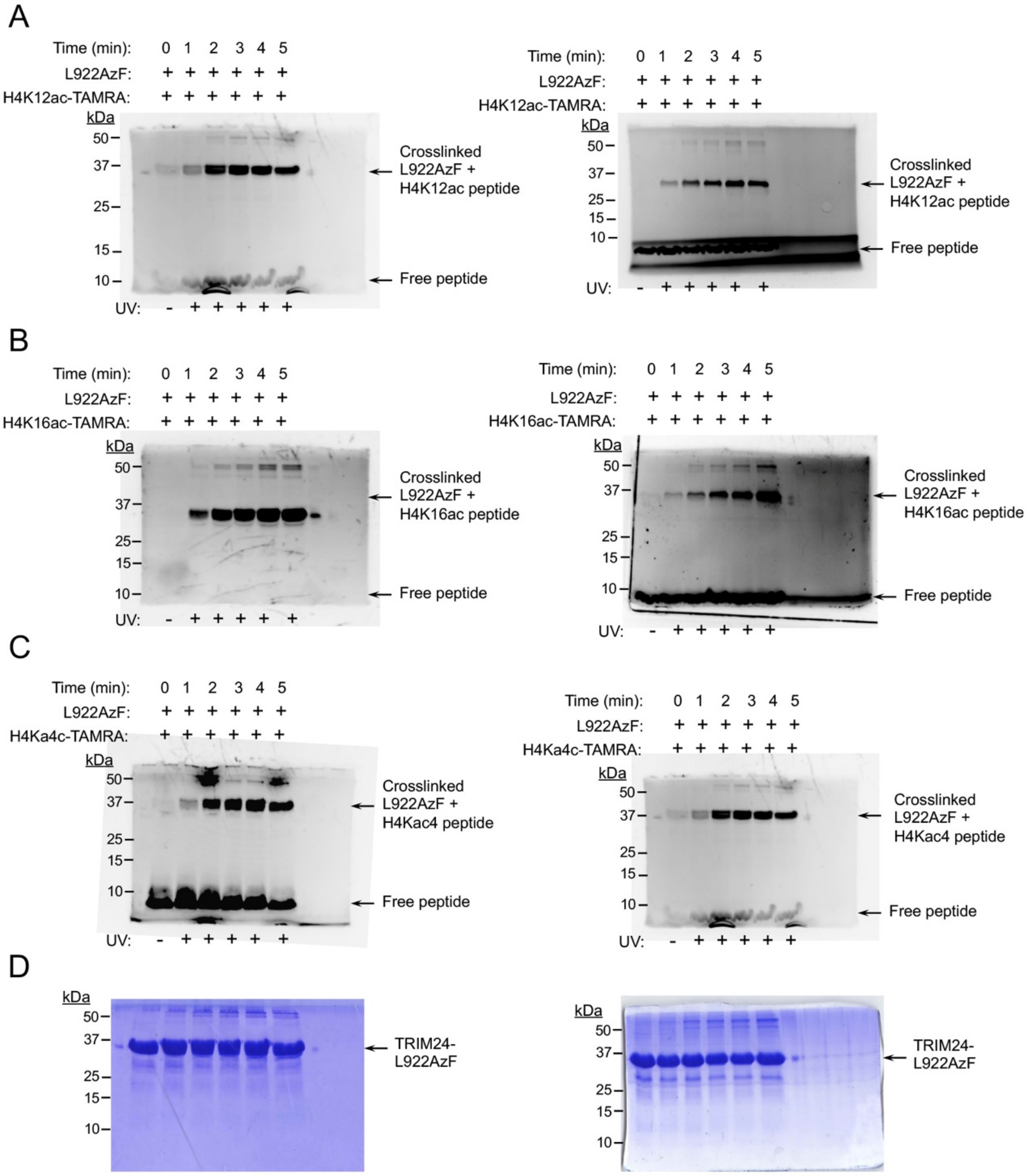


**Figure S2.** Photocrosslinking of engineered TRIM24-L922AzF with acetylated histone H4 peptides. The engineered TRIM24 bromodomain containing 4-azido-L-phenylalanine (AzF) at position L922 was incubated with TAMRA-labelled histone H4 peptides and subjected to UV irradiation at 365 nm for the indicated times. (A-C) In-gel fluorescence analysis showing time-dependent photo-crosslinking of TRIM24-L922AzF with TAMRA-labelled H4K12ac (A), H4K16ac (B), and H4Kac4 (K5ac, K8ac, K12ac, and K16ac) (C) peptides. The crosslinked TRIM24-L922AzF-H4 peptide complexes are observed at the expected higher molecular mass, while unreacted/free TAMRA-labelled peptides migrate near the bottom of the gel. Each crosslinking experiment was performed in duplicate. (D) Representative Coomassie-stained SDS-PAGE gels confirming the presence of TRIM24-L922AzF protein in the corresponding samples.

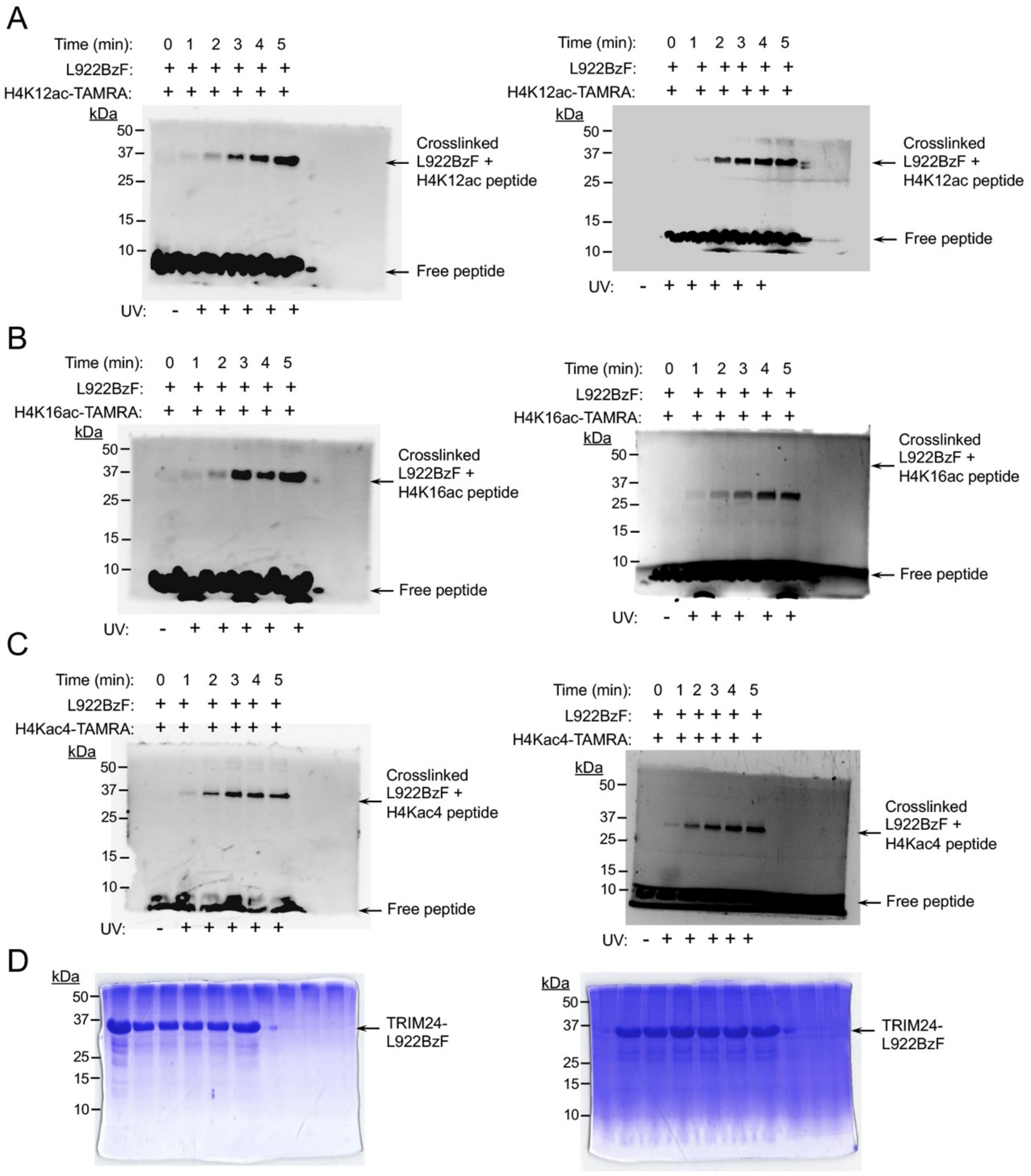


**Figure S3.** Photocrosslinking of engineered TRIM24-L922BzF with acetylated histone H4 peptides. The engineered TRIM24 bromodomain containing 4-azido-L-phenylalanine (AzF) at position L922 was incubated with TAMRA-labelled histone H4 peptides and subjected to UV irradiation at 365 nm for the indicated times. (A-C) In-gel fluorescence analysis showing time-dependent photo-crosslinking of TRIM24-L922BzF with TAMRA-labelled H4K12ac (A), H4K16ac (B), and H4Kac4 (K5ac, K8ac, K12ac, and K16ac) (C) peptides. The crosslinked TRIM24-L922BzF-H4ac peptide complexes are observed at the expected higher molecular mass, while unreacted/free TAMRA-labelled peptides migrate near the bottom of the gel. Each crosslinking experiment was performed in duplicate. (D) Representative Coomassie-stained SDS-PAGE gels confirming the presence of TRIM24-L922BzF protein in the corresponding samples.

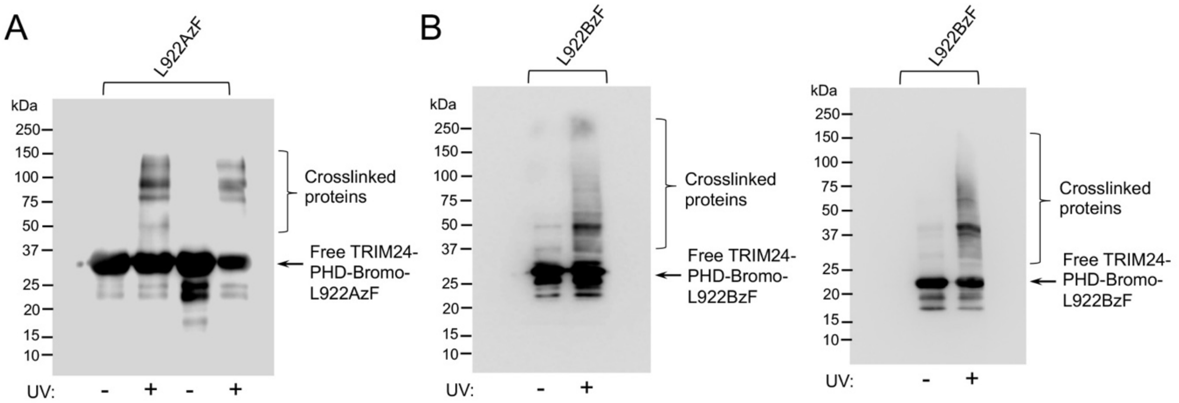


**Figure S4.** Photocrosslinking of TRIM24 bromodomain mutants with cellular proteins. HEK293T cells were treated with the deacetylase inhibitor trichostatin A (TSA) to increase cellular protein acetylation. Hyperacetylated cell lysates were incubated with purified TRIM24 PHD–bromodomain L922AzF or L922BzF mutant proteins and subjected to 365-nm UV irradiation to induce photocrosslinking. Crosslinked products were analyzed by immunoblotting using an anti-6×His antibody. UV-dependent higher-molecular-weight species containing the His-tagged TRIM24 PHD–bromodomain mutants are indicated as crosslinked proteins, whereas the corresponding bands of the free TRIM24 PHD–bromodomain mutants are indicated as Free L922AzF or Free L922BzF. (A, B) Uncropped Western blot images corresponding to the photocrosslinking experiments shown in Figures 6B and 7B, respectively. (A) L922AzF and (B) L922BzF.

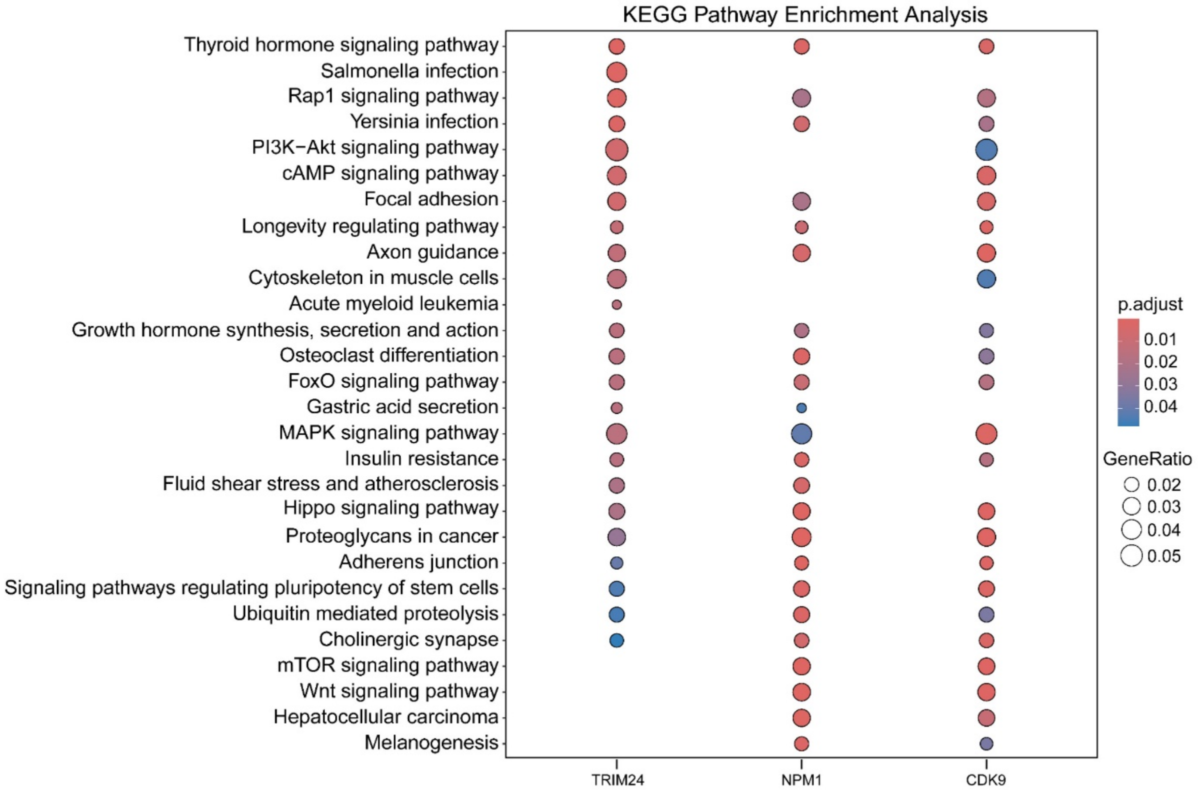


**Figure S5.** KEGG pathway enrichment analysis of TRIM24, CDK9, and NPM1. Dot plot showing significantly enriched KEGG pathways associated with TRIM24, CDK9, and NPM1. Dot size represents the GeneRatio, indicating the proportion of genes associated with each pathway, while dot color represents the adjusted p-value (p.adjust), with red indicating greater statistical significance and blue indicating lower statistical significance. The x-axis represents the proteins/genes analyzed, and the y-axis represents the enriched KEGG pathways.

**HPLC REPORT**

| | | |
|---|---|---|
| Sample: | Pep-694 ARTKQTARKSTGGKAPRKQLAT(Kac)AARKSAPATG | Analyzed date: 24-03-2023 |
| Analyst: | Dr.RS-SBio | |
| Column: | 4.6×250mm,Sinochrom ODS-BP 5μm | |
| Solvent A | A: 0.1% Trifluoroacetic Acid in 100% Acetonitrile | |
| Solvent B | B: 0.1% Trifluoroacetic Acid in 100% Water | |

Gradient:

| | A | B |
|---|---|---|
| 0.0min | 10% | 90% |
| 25.0min | 35% | 65% |
| 25.1min | 100% | 0% |
| 30.0min | | Stop |

Volume: 5μl

Wavelength: 220nm

Flow rate: 1.0ml/min

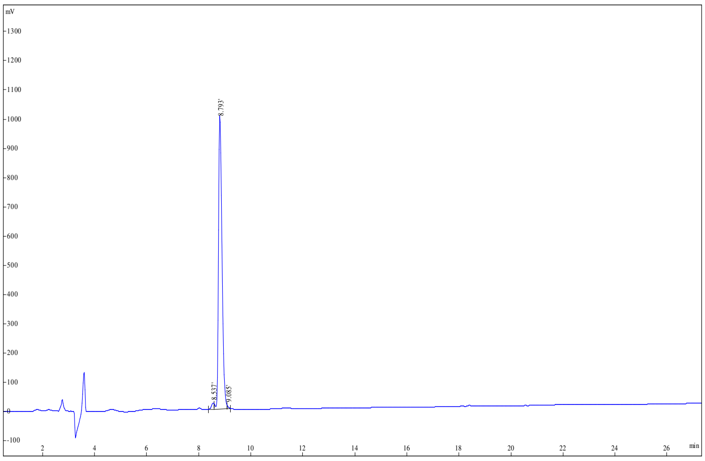

| Rank | Time | Area | Conc. |
|---|---|---|---|
| 1 | 8.537 | 150619 | 1.553 |
| 2 | 8.793 | 9519529 | 98.16 |
| 3 | 9.085 | 27391 | 0.2825 |
| Total | | | 100 |

**Figure S6.** HPLC purity traces for the H3(1-33)K23ac peptide.

**MASS SPECTROMETRY REPORT**

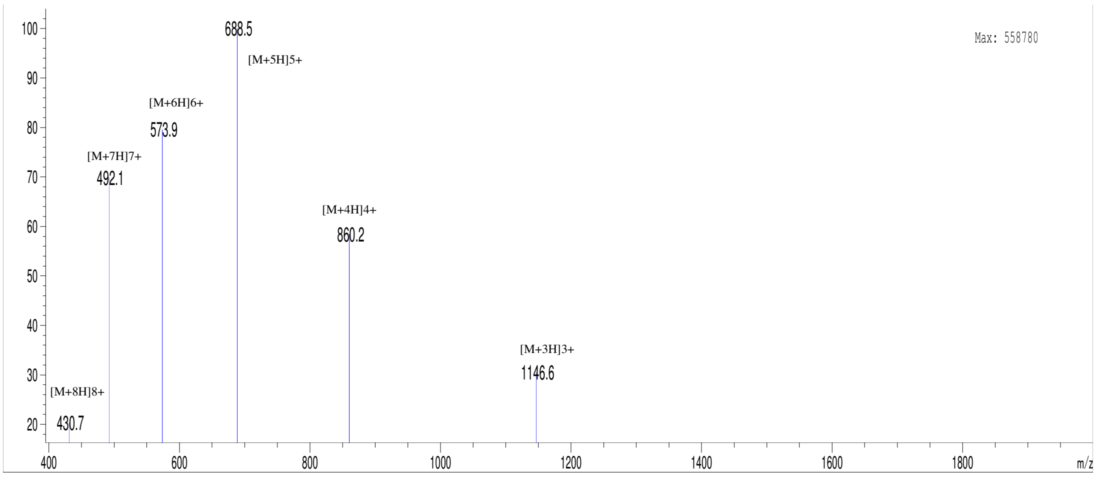


Sample Description

| | | | | | |
|---|---|---|---|---|---|
| | | Instrument | Agilent-6125B | | |
| Analyzed date: | 25-03-2023 | Probe: | ESI | Probe Bias: | +4.5kv |
| Analyst: | Dr.AR-SBio | Nebulizer Gas Flow: | 1.5L/min | Detector: | 1.5kv |
| Sample: | Pep-694 ARTKQTARKSTGGKAPRKQLAT(Kac)AARKSAPATG | CDL: | -20.0v | T. Flow: | 0.2ml/min |
| M.W.: | 3436.92 | CDL Temp.: | 250 °C | B. Conc.: | 50%H2O/50%ACN |

**Figure S7.** MS spectra for the H3(1-33)K23ac peptide.

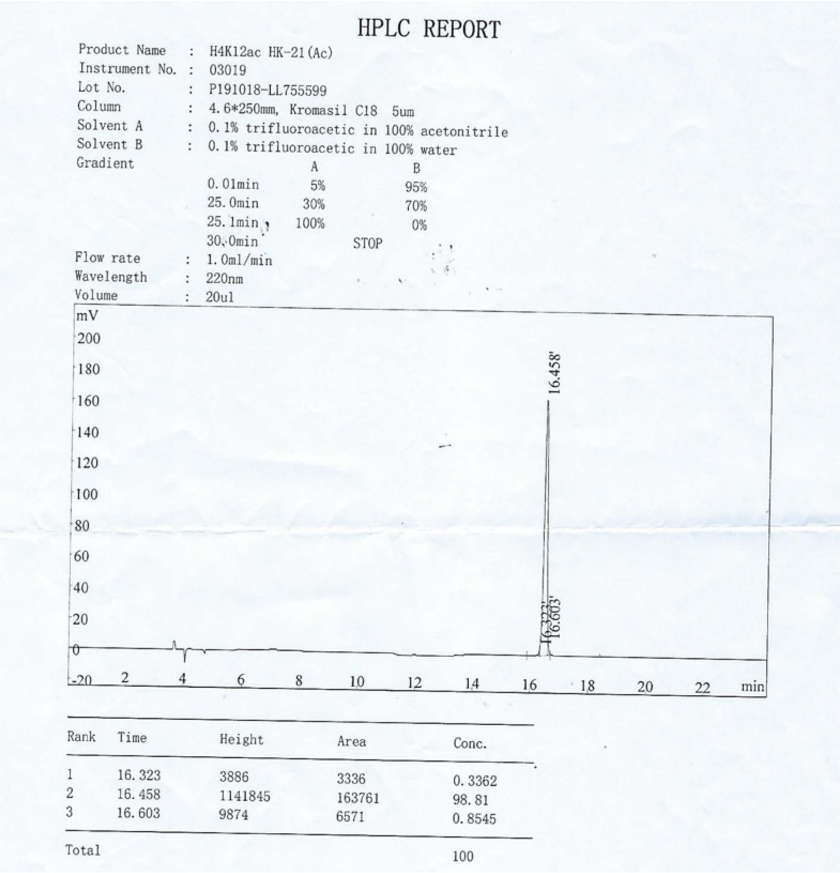

HPLC REPORT

Product Name : H4K12ac HK-21(Ac)
Instrument No. : 03019
Lot No. : P191018-LL755599
Column : 4.6*250mm, Kromasil C18 5um
Solvent A : 0.1% trifluoroacetic in 100% acetonitrile
Solvent B : 0.1% trifluoroacetic in 100% water
Gradient

| | A | B |
|---|---|---|
| 0.01min | 5% | 95% |
| 25.0min | 30% | 70% |
| 25.1min | 100% | 0% |
| 30.0min | STOP | |

Flow rate : 1.0ml/min
Wavelength : 220nm
Volume : 20ul



| Rank | Time | Height | Area | Conc. |
|---|---|---|---|---|
| 1 | 16.323 | 3886 | 3336 | 0.3362 |
| 2 | 16.458 | 1141845 | 163761 | 98.81 |
| 3 | 16.603 | 9874 | 6571 | 0.8545 |
| Total | | | | 100 |

**Figure S8.** HPLC purity traces for the H4(1-20)K12ac peptide.

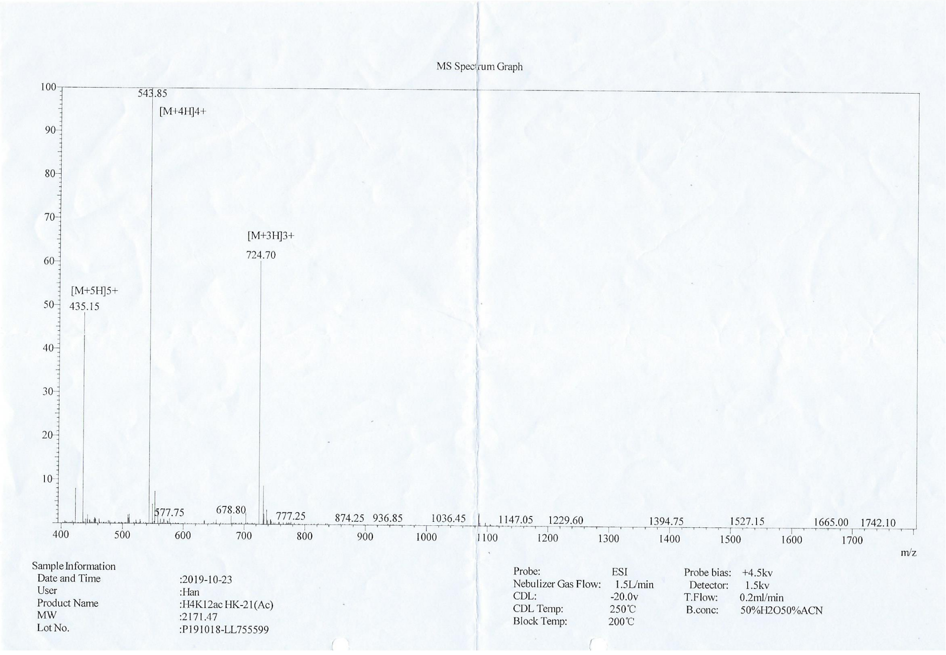


**Figure S9.** Mass spectra for the H4(1-20)K12ac peptide.

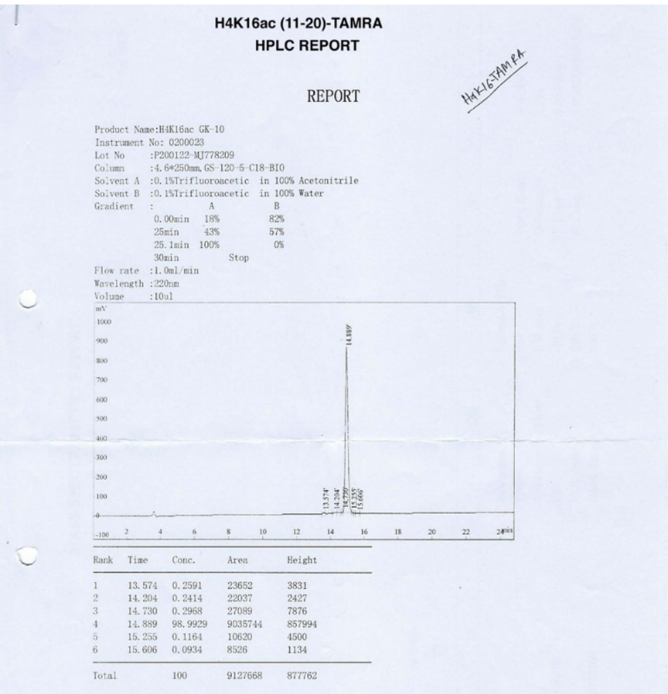

**H4K16ac (11-20)-TAMRA**

**HPLC REPORT**

REPORT

Product Name:H4K16ac GK-10
Instrument No: 0200023
Lot No :P200122-MJ778209
Column :4.6*250mm, GS-120-5-C18-BIO
Solvent A :0.1%Trifluoroacetic in 100% Acetonitrile
Solvent B :0.1%Trifluoroacetic in 100% Water

| Gradient | A | B |
|---|---|---|
| 0.00min | 18% | 82% |
| 25min | 43% | 57% |
| 25.1min | 100% | 0% |
| 30min | Stop | |

Flow rate :1.0ml/min
Wavelength :220nm
Volume :10ul



| Rank | Time | Conc. | Area | Height |
|---|---|---|---|---|
| 1 | 13.574 | 0.2591 | 23652 | 3831 |
| 2 | 14.204 | 0.2414 | 22037 | 2427 |
| 3 | 14.730 | 0.2968 | 27089 | 7876 |
| 4 | 14.889 | 98.9929 | 9035744 | 857994 |
| 5 | 15.255 | 0.1164 | 10620 | 4500 |
| 6 | 15.606 | 0.0934 | 8526 | 1134 |
| Total | | 100 | 9127668 | 877762 |

Figure S10. HPLC purity traces for the H4(11-20)K16ac peptide.

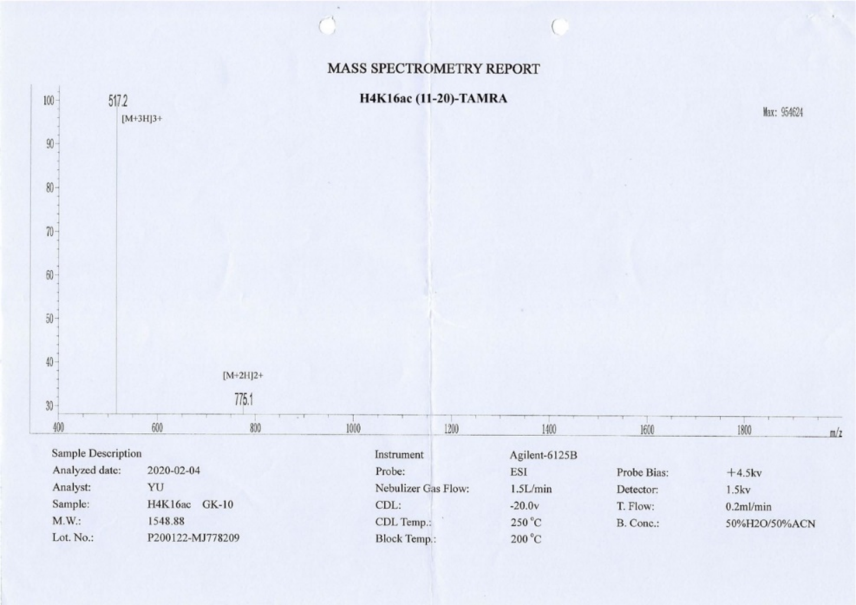

**MASS SPECTROMETRY REPORT**

**H4K16ac (11-20)-TAMRA**



| | | | | | |
|---|---|---|---|---|---|
| Sample Description | | Instrument | Agilent-6125B | | |
| Analyzed date: | 2020-02-04 | Probe: | ESI | Probe Bias: | +4.5kv |
| Analyst: | YU | Nebulizer Gas Flow: | 1.5L/min | Detector: | 1.5kv |
| Sample: | H4K16ac GK-10 | CDL: | -20.0v | T. Flow: | 0.2ml/min |
| M.W.: | 1548.88 | CDL Temp.: | 250 °C | B. Conc.: | 50%H2O/50%ACN |
| Lot. No.: | P200122-MJ778209 | Block Temp.: | 200 °C | | |

**Figure S11.** Mass spectra for the H4(11-20)K16ac peptide.

**H4K5acK8acK12acK16ac (1-20)-TAMRA**

**HPLC REPORT**

| | | | |
|---|---|---|---|
| Sample: | Pep-169 SGRGK(ac)GGK(ac)GLGK(ac)GGAK(ac)RHRK-TAMRA | | Analyzed date: 08-11-2020 |
| Analyst: | Dr.RS-SBio | | |
| Column: | Symmetrix ODS-R, 4.6*250mm, 5μm | | |
| Solvent A | A: 0.1% Trifluoroacetic Acid in 100% Acetonitrile | | |
| Solvent B | B: 0.1% Trifluoroacetic Acid in 100% Water | | |
| Gradient: | | A | B |
| | 0.0min | 18% | 82% |
| | 25.0min | 43% | 57% |
| | 25.1min | 100% | 0% |
| | 30.0min | Stop | |
| Volume: | 20μl | | |
| Wavelength: | 220nm | | |
| Flow rate: | 1.0ml/min | | |

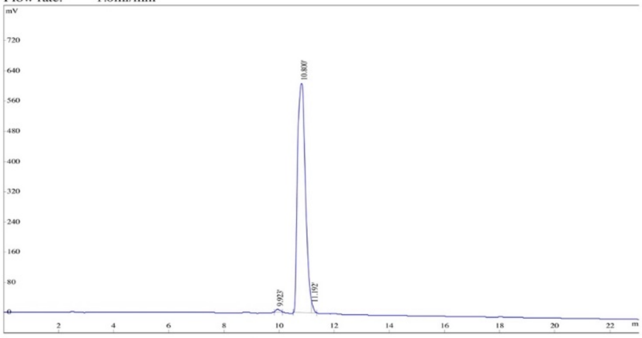


| Rank | Time | Conc. | Area | Height |
|---|---|---|---|---|
| 1 | 9.923 | 0.7119 | 83545 | 7885 |
| 2 | 10.800 | 98.4844 | 11557687 | 608544 |
| 3 | 11.192 | 0.8037 | 94316 | 21570 |
| Total | | 100 | 11735548 | 637999 |

**Figure S12.** HPLC purity traces for the H4(1-20)Kac4 (K5/K8/K12/K16) peptide.

**H4K5acK8acK12acK16ac (1-20)-TAMRA**

**MASS SPECTROMETRY REPORT**

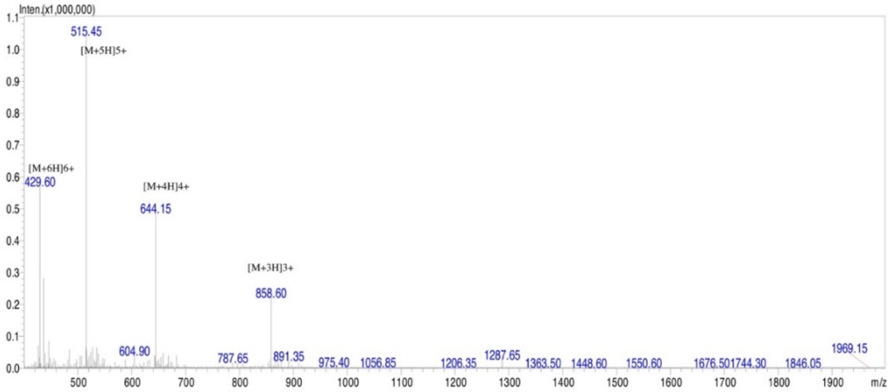


| | | | | | |
|---|---|---|---|---|---|
| Sample Description | | Instrument | SHIMADZU LCMS-2020 | | |
| Analyzed date: | 09-11-2020 | Probe: | ESI | Probe Bias: | 4.5kv |
| Analyst: | Dr.AR-SBio | Nebulizer Gas Flow: | 1.5L/min | Detector: | 1.5kv |
| Sample: | Pep-169 SGRGK(ac)GGK(ac)GLGK(ac)GGAK(ac)RHRK-TAMRA | CDL: | -20.0v | T. Flow: | 0.2ml/min |
| M.W.: | 2572.89 | CDL Temp.: | 250 °C | B. Conc.: | 50%H2O/50%ACN |

**Figure S13.** MS spectra for the H4(1-20)Kac4 (K5/K8/K12/K16) peptide.
